# Persistence of vestibular function in the absence of glutamatergic transmission from hair cells

**DOI:** 10.1101/2025.10.15.682551

**Authors:** Mohona Mukhopadhyay, Ruchi Modgekar, Aizhen Yang-Hood, Kevin K. Ohlemiller, Valentin Militchin, Maolei Xiao, Zhijun Shen, Nicholas R Rensing, Michael Wong, Suh Jin Lee, Rebecca P. Seal, Mark E. Warchol, Susan E. Maloney, Carla M. Yuede, Mark A. Rutherford, Tina Pangrsic

## Abstract

Quantal synaptic transmission in vestibular end-organs is glutamatergic. Although genetic deletion of *Slc17a8* (termed *Vglut3*) leads to deafness in mice, the dependence of vestibular function on VGLUT3-mediated quantal transmission is unknown. Here, we investigated the vestibular phenotype of *Vglut3^-/-^* mice at the cellular, systems, and behavioral levels. The type-II vestibular hair cells (VHCs) in *Vglut3^+/+^*mice were strongly immunoreactive for VGLUT3, while type-I VHCs showed poor immunoreactivity. In *Vglut3^-/-^* mice quantal synaptic transmission in utricular calyces was reduced in rate and amplitude by > 95%. *In vivo* recordings of spontaneous activity in the vestibular nerve revealed similar action potential rates and regularity in *Vglut3^+/+^*and *Vglut3^-/-^* mice, suggesting a divergent underlying mechanism compared to the silent *Vglut3^-/-^* auditory nerve. In behavioral studies, *Vglut3^-/-^* mice did not exhibit considerable sensorimotor or balance deficits. Collectively, these data support the view that non-quantal transmission is the predominant mode of neurotransmission between type I VHCs and vestibular calyceal afferent neurons. We propose that non-quantal transmission alone underlies the apparently normal vestibular nerve physiology and behavioral function in *Vglut3^-/-^* mice.

## Introduction

A faithfully choreographed interaction between the peripheral vestibular system and the central nervous system is how mammals perceive head motion and maintain balance in their daily lives. The organs constituting the peripheral vestibular system in amniotes, the otoliths organs i.e., utricle and saccule, and the three semicircular canals, execute these functions owing to a precise interplay between various modes of synaptic transmission.

The calyx evolved in amniotes as a specialized postsynaptic terminal to the presynaptic type I hair cell in the vestibular organs (Eatock & Songer, 2011). Several reports have now suggested various mechanisms of neurotransmission in the calyx, including but not limited to, quantal transmission and faster nonquantal transmission (reviewed in Mukhopadhyay & Pangrsic, 2022). While the former directly depends on vesicular exocytosis of the neurotransmitter glutamate, the latter may involve factors such as cleft-potential, resistive coupling, or potassium concentration, and may be influenced by proton and glutamate accumulation in the cleft (Govindaraju *et al*, 2023; Contini *et al*, 2012, 2020, 2017; Spaiardi *et al*, 2020; Lim *et al*, 2011; Highstein *et al*, 2014; Sadeghi *et al*, 2014; Contini *et al*, 2024).

Vesicular glutamate transporters (VGLUT) constitute a family of proteins responsible for packaging the excitatory neurotransmitter glutamate into secretory vesicles and thus represent vital components of quantal neurotransmission at excitatory synapses (reviewed in Pietrancosta *et al*, 2020; El Mestikawy *et al*, 2011). Exocytosis of vesicular glutamate activates post-synaptic glutamate receptors, thus depolarizing the vestibular fibers of the VIII^th^ nerve (Sadeghi *et al*, 2014).

Three genes *SLC17A6, SLC17A7* and *SLC17A8* encode three related VGLUT proteins (Fremeau *et al*, 2004). While VGLUT1 and VGLUT2 are the predominant isoforms in the mammalian nervous system (Fremeau *et al*, 2004), VGLUT3 is more restricted (Seal & Edwards, 2006). Cochlear inner hair cells (IHC) rely on VGLUT3 for sound-evoked glutamate release (Seal *et al*, 2008; Ruel *et al*, 2008). Genetic disruption of *Vglut3* in mice abolishes glutamate release and postsynaptic activity at cochlear IHC afferent synapses resulting in deafness (Seal *et al*, 2008). *Vglut3^-/-^* mice also exhibit progressive degeneration of ribbon synapses and spiral ganglion neurons in the absence of glutamate release (Seal *et al*, 2008; Kim *et al*, 2019). A non-syndromic autosomal dominant deafness in humans, DFNA25, is associated with sequence variants in *SLC17A8* (Ruel *et al*, 2008; Joshi *et al*, 2021; Ryu *et al*, 2016). In zebrafish, the removal of Vglut3 function leads to loss of acoustic startle reflex and vestibulo-ocular reflex (Obholzer *et al*, 2008).

Although germline *Vglut3^-/-^* mice suffer from congenital deafness and, on some genetic backgrounds, infrequent nonconvulsive seizures, no overt symptoms of vestibular dysfunction have been reported (Seal *et al*, 2008; Regalado Núñez *et al*, 2024). VGLUT3 expression has been detected in hair cells in the different vestibular organs (e.g. Schraven *et al*, 2012; Zhang *et al*, 2011). However, detailed studies on the localization of this family of transporters within different vestibular organs, zones, and cell types in mice are not available. Quantal transmission at the synapses between type-I VHCs and the postsynaptic calyces of vestibular ganglion neurons has been shown to be glutamatergic (Bonsacquet *et al*, 2006; Contini *et al*, 2017; Demêmes *et al*, 1995; Dulon *et al*, 2009; Highstein *et al*, 2015; Holt *et al*, 2007; Kirk *et al*, 2017; Sadeghi *et al*, 2014; reviewed in Mukhopadhyay & Pangrsic, 2022). However, the specific roles of VGLUT1-3 in glutamatergic neurotransmission at these specialized calyceal synapses remains unclear.

In the present study we addressed these questions by performing a series of immunohistochemistry, RNAscope, *in vivo-* and *ex vivo-* cell physiology experiments and behavioral studies. Our results suggest that VGLUT3 is essential for quantal transmission at peripheral vestibular afferent synapses, and that its loss may be largely compensated for by mechanisms of nonquantal transmission to support adapted vestibular functions in *Vglut3^-/-^*mice.

## Results

### *Vglut3^-/-^* mice with profound hearing loss show no obvious signs of disequilibrium

Loss of VGLUT3 from cochlear hair cells results in deafness due to the absence of afferent transmission (Ruel *et al*, 2008; Seal *et al*, 2008; Kim *et al*, 2019; Akil *et al*, 2012), and some *SLC17A8* variants are associated with progressive hearing loss in humans (Joshi *et al*, 2021; Ruel *et al*, 2008; Ryu *et al*, 2016). In the same cohorts of mice subjected to tests of balance (**Fig. 1** and **Suppl. Fig 1**), we first checked the acoustic startle response (ASR) in response to white noise bursts, which has been shown to decrease with age in B6 mice (Shoji *et al*, 2016). We compared *Vglut3^+/+^* and *Vglut3^-/-^* littermates at 3, 8, and 12 months of age. At 3-months, all *Vglut3^+/+^* mice had large startle responses that grew with the acoustic stimulus, while *Vglut3^-/-^* littermates, on average, did not (**Fig. 1A-B**; **Table 1**). At 8 and 12 months, ASR magnitudes in *Vglut3^+/+^*were much smaller than at 3 months, presumably due to the *Cdh23*^c.753A^ age-related hearing loss phenotype on the C57BL/6 background (Johnson *et al*, 2017), since the normal-hearing CBA/CaJ mice do not show age-related reduction in ASR (Ison & Allen, 2003). Still, compared to *Vglut3^-/-^*, there was a main effect of genotype and an interaction between genotype and dB SPL (**Table 1**). In *Vglut3^+/+^,* all mice significantly startled at 3 and 8 months, but, consistent with age-related hearing loss, only 6 of 8 mice startled at 12 months. Interestingly, at the highest stimulus level of 120 dB SPL, some *Vglut3^-/-^* mice did startle significantly in the startle trials compared to the no-stimulus trials (**Fig. 1B-C**; 3 of 10 at 3 months, 2 of 11 at 8 months, 1 of 11 at 12 months; paired *t*-test, *p* < 0.05), possibly due to saccular activation (Jones *et al*, 2010).

**Figure 1.**
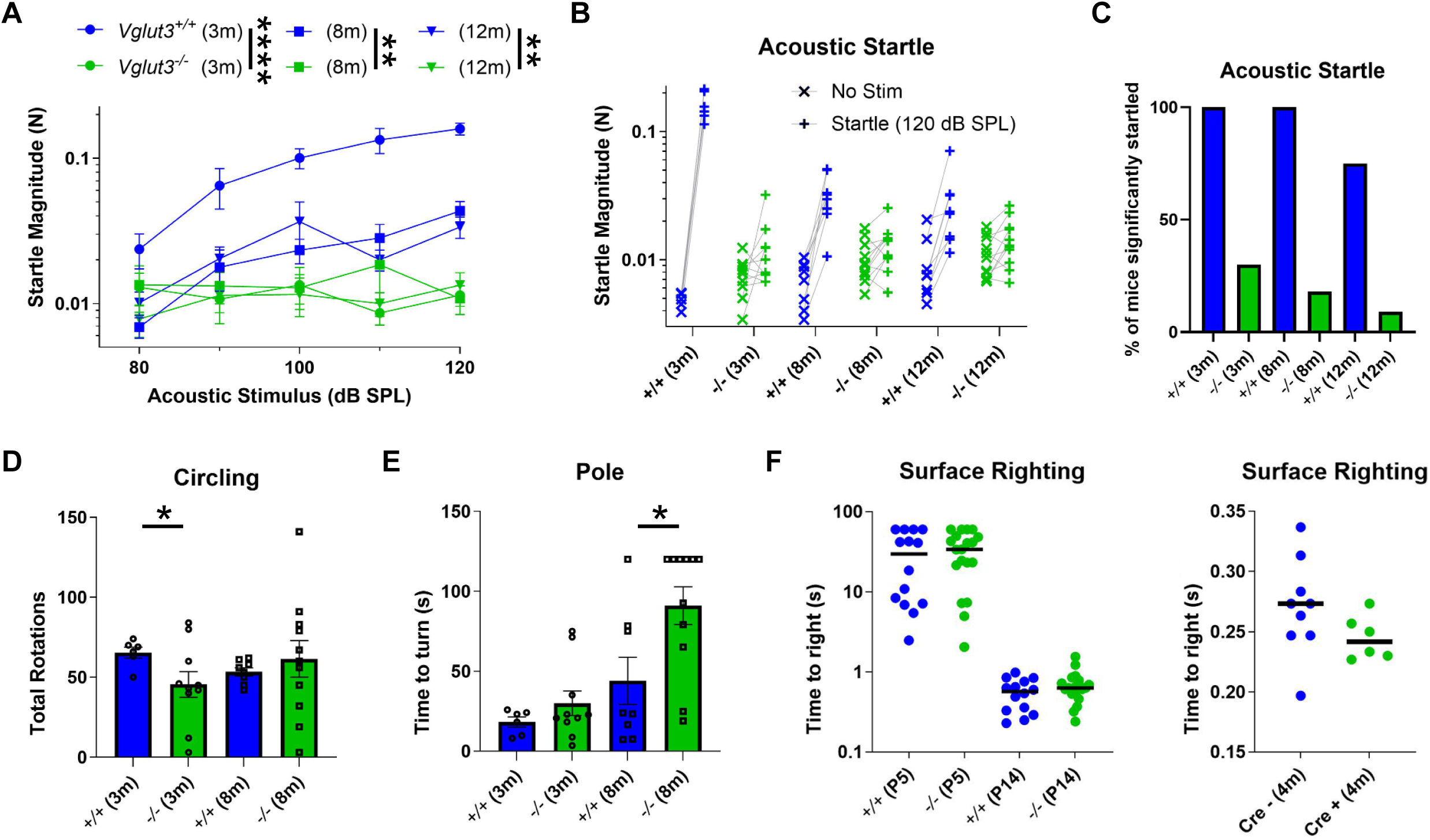
Tests of acoustic startle and vestibular equilibrium. **A)** Growth of acoustic startle response magnitude with increasing stimulus level was significantly greater in *Vglut3*^+/+^ mice compared to *Vglut3*^-/-^ (see Table 1). **B)** Mean startle magnitude for no stimulation trials (x) and startle trials at 120 dB SPL (+), connected by a line for each mouse tested. **C)** Percentage of mice significantly startled decreased with age (n = 10 trial pairs per mouse). **D)** Circling behaviour had significantly more variance in the *Vglut3*^-/-^ group of mice compared to *Vglut3*^+/+^ at 3 (*p* = 0.032) but not 8 (*p* = 0.3) months of age (Kolmogorov-Smirnov test). **E)** At 8 months, the *Vglut3*^-/-^ mice took longer to turn and climb down the pole (*p* = 0.023, Mann Whitney U test). **F)** There was no difference in the surface righting reflex time in juvenile (P5) or young (P14) *Vglut3*^+/+^ mice compared to *Vglut3*^-/-^ (*left*). Similarly, surface righting did not appear impaired in *Vglut3-*conditional KO mice at 4 months of age (*right*).

**Table 1.** Statistical comparison of ASR magnitudes in *Vglut3^+/+^* and *Vglut3^-/-^*littermates at 3, 8, and 12 months of age. The values in the table represent the *p*-values of the 2-way repeated measures ANOVA with Bonferroni corrected multiple comparisons.

| Age (in months) | Genotype effect | Genotype × dB SPL interaction |
| --- | --- | --- |
| 3 | $< 10^{-4}$ | $< 10^{-4}$ |
| 8 | 0.0065 | 0.013 |
| 12 | 0.0063 | 0.038 |

VGLUT3 is expressed in vestibular end-organs as well (Schraven *et al*, 2012; Zhang *et al*, 2011), suggesting it may be responsible for glutamatergic synaptic transmission in the inner ear, and could underlie behaviors that rely, in part, on vestibular function. *Vglut3^-/-^* and *Vglut3^+/+^* mice had similar body weight (**Suppl. Fig. 1A**) and *Vglut3^-/-^* mice did not exhibit an increase in circling behavior, on average (**Fig. 1D**), a phenotype often associated with pathology of the vestibular sensory epithelium (Bloom & Hultcrantz, 1994). However, unlike the means, the variance in number of total rotations was much greater in the *Vglut3^-/-^* mice (3 months: range 3-84; 8 months: range 3-141) compared with *Vglut3^+/+^*mice (3 month: range 50-74; 8 months: 50-62), and the shapes of the two distributions were significantly different at 3 but not at 8 months (**Fig. 1D**). In the screen test, a test of climbing at inclined angles, there were no significant differences between genotypes for the 60°, 90,° or inverted screen at 3 or 8 months (**Suppl. Fig. 1B**). In the stationary rod, constant speed rotarod, and accelerating rotarod tests, there were no significant overall effects of genotype and no interactions between genotype and trial number (**Suppl. Fig. 1D**). In contrast, in the pole test, *Vglut3*^-/-^ mice took significantly longer to turn and face downward to walk down the pole at 8 months (**Fig. 1E**). The *Vglut3*^-/-^ mice displayed no difficulties swimming, in fact, they swam faster and further than *Vglut3*^+/+^ at 8 months of age (**Suppl. Fig. 1C**). Moreover, they showed normal surface righting reflexes at P5 and P14 (**Fig. 1F**) and normal air righting reflex at P18 (% of mice that fully righted: 79% WT; 75% HET; 80% KO; Chi-Square test). To address the possibility that righting reflexes in the germline *Vglut3^-/-^* mice were similar to *Vglut3^+/+^* because they adapted early in postnatal development to compensate with other sensory modalities, we allowed *Vglut3-*conditional KO mice to mature before administering tamoxifen (1 mg per day for 5 days) at 3 months. Although they were confirmed deaf by auditory brainstem response at 3.5 months (not shown), the Cre+ mice displayed similar surface righting reflex performance to Cre– mice at 4 months (**Fig. 1F**) and similar air righting (78% Cre-; 69% Cre+; Chi-Square test).

*Vglut3*^-/-^ and *Vglut3^+/+^* control mice underwent video-EEG monitoring starting at 6 months of age for two weeks continuously. *Vglut3*^-/-^ mice had normal awake and asleep background EEG patterns that were indistinguishable from control mice (**Suppl. Fig. 1E**), with predominantly low amplitude, higher frequency theta activity while awake and higher amplitude, lower frequency delta activity during non-REM sleep. In contrast to previous reports (Seal *et al*, 2008), no abnormal spike and wave discharges or electrographic seizures were detected in either *Vglut3*^-/-^ or control mice, possibly suggesting mouse strain-dependent manifestations of the phenotype.

In summary, although the ASR was absent or extremely deficient in *Vglut3*^-/-^ mice, no major and consistent differences across age groups were observed in the battery of open-field and sensorimotor performance tests. The *Vglut3*^-/-^ mice exhibited greater variance in circling behavior at 3 months, took longer to turn and climb down the pole or did not attempt to climb down at 8 months, and swam faster and further at 8 months. Similarly, previous study reported that some *Vglut3*^-/-^ mice took longer to cross an easy beam (Regalado Núñez *et al*, 2024). These mildly altered behaviors could possibly result from impairment of circuits not related specifically to vestibular sensation but to increased anxiety due to the lack of VGLUT3 from inhibitory neurons in the brain (Amilhon *et al*, 2010).

### *Vglut* transcript and protein expression in vestibular hair cells and ganglion neurons

In the cochlea, inner hair cells rely exclusively on VGLUT3 (Seal *et al*, 2008), while in outer hair cells there is a minor contribution by VGLUT2 to glutamatergic transmission, as well (Weisz *et al*, 2021). Auditory nerve fibers are thought to express predominantly VGLUT1, at least in guinea pig (Zhou *et al*, 2007). Vestibular type II hair cells, but not type I hair cells, have been reported to express VGLUT3 (Schraven *et al*, 2012). Vestibular ganglion neurons (VGN) in rat are reported to express VGLUT1 and/or VGLUT2 (Wang *et al*, 2007; Zhang *et al*, 2011). Although hair cells and VIIIth nerve are generally considered to express VGLUT3 and VGLUT1/2, respectively, some studies have reported VGLUT3 in afferent axons in vestibular epithelia and VGLUT1 in VHCs (Wang *et al*, 2007; Zhang *et al*, 2011). Given the tight juxtaposition of hair cells and neurons in the vestibular end-organs, particularly the type I hair cell to calyx synapse, and depending on the resolution and sensitivity of the microscopy immuno-detection used, close apposition could appear as co-localization and low-level expression could appear as absence. Here, we addressed this issue anatomically at the RNA and protein level with high-resolution confocal and Airyscan microscopy in histological sections.

Otolith organs and cristae were probed for *Vglut1* and *Vglut2* mRNA levels and immuno-stained for Myosin7a and a combination of neuron-specific beta-III-tubulin (TuJ1) and neurofilament H (Nf-H) to label hair cells and vestibular neurons, respectively (**Fig. 2**, **Suppl. Fig. 2**). *Vglut1* and *2* mRNA puncta in the sensory epithelium mostly appeared as a diffuse but non-homogeneous layer of fluorescence. When present in spots, rarely more than three distinct spots were observed per hair cell (or in its direct vicinity) in any of the vestibular peripheral organs (**Fig. 2A-D**). On the contrary, vestibular ganglion neurons in the same cryosections hosted many such spots, indicating good staining quality and significant mRNA expression (**Fig. 2E-H**). Typically, the cells are considered to be positive for a particular RNAscope probe when at least three such spots are observed per cell. Although it is difficult to say with certainty, we consider such diffused staining pattern and low abundance of spots in the hair cells as a sign of very low expression of the *Vglut1* and *Vglut2*, or background, in particular in comparison with the results obtained from the vestibular neurons on the same microscope slides (**Fig. 2E-H**). We also note that the sparse mRNA signals in the hair cell epithelium could be located in the neuronal postsynaptic compartments. In any case, we observed no obvious differences in the staining pattern or abundance of *Vglut1* or *Vglut2* in the region of the vestibular hair cells of *Vglut3*^+/+^ as compared to *Vglut3*^-/-^ mice, suggesting no upregulation of either VGLUT in the absence of VGLUT3.

**Figure 2.**
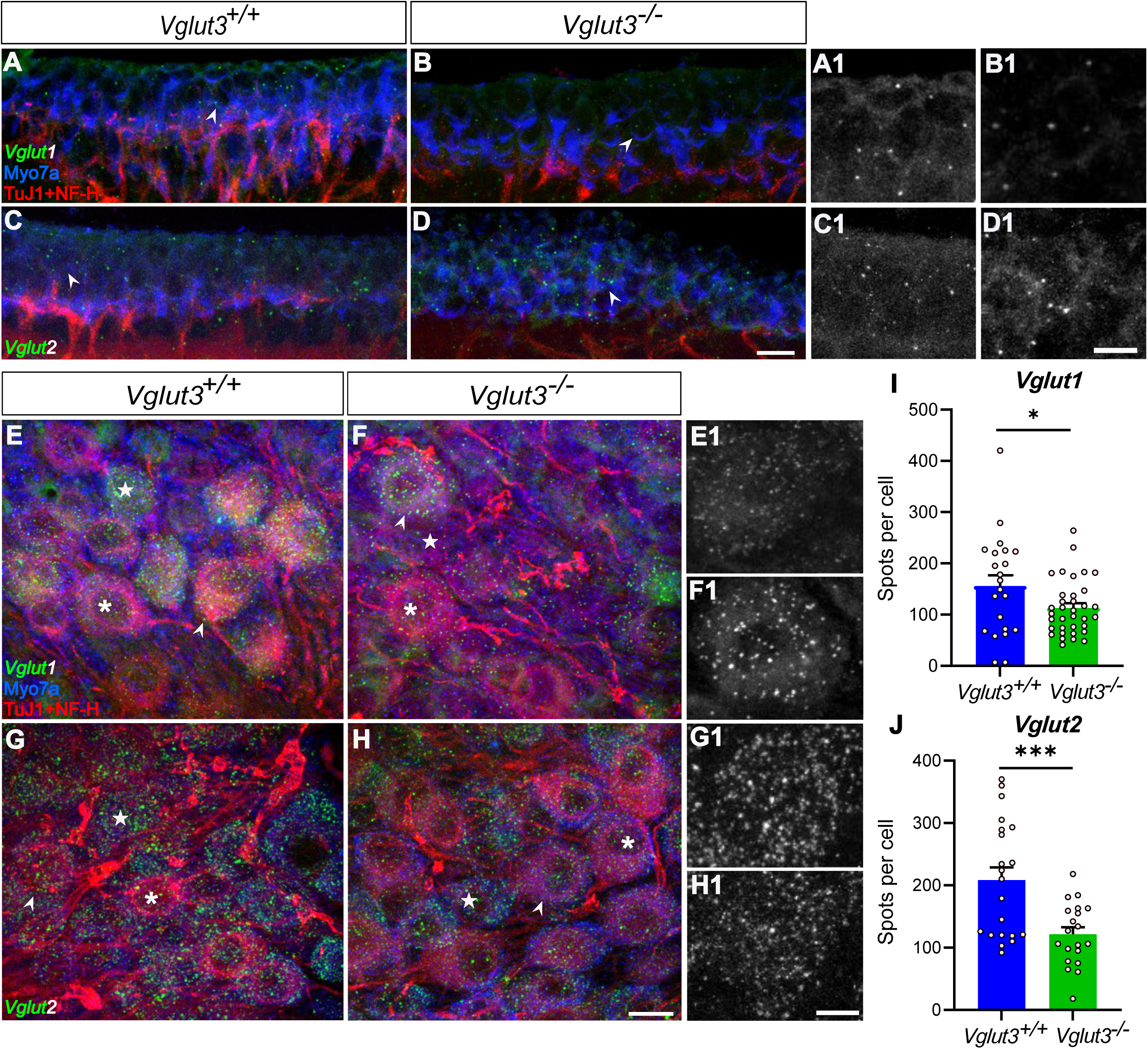
*Vglut1* and *2* transcripts in the vestibular ganglion and sensory epithelia. **A-H)** *Vglut1* and *Vglut2* mRNA transcripts in the utriculi (A-D) and vestibular ganglion (E-H) of 12-week-old *Vglut3*^+/+^ and *Vglut3*^-/-^ mice. Note a mostly diffuse staining pattern with only a few distinct spots for both mRNA probes (*green*) in the utriculi, but, several distinct spots in vestibular neurons. Antibodies against myosin VIIa (Myo7a) and a combination of neurofilament-H (NF-H) and beta-tubulin III (TuJ1) were used to identify VHCs and vestibular neurons, respectively. Scale bars: A-D:5 µm, A1-D1: 3 µm, E-H: 2 µm, E1-H1: 1 µm. Arrowheads point at the regions displayed at high magnification in the inserts A1-D1 and E1-H1 (the signal from the respective mRNA probes only). Examples of NF-H/Tuj1-rich (*asterisks*) and –poor (*stars*) neurons. **I-J)** Average abundance of *Vglut1* and *Vglut2* transcripts in the vestibular neurons of *Vglut3*^+/+^ and *Vglut3*^-/-^ mice. Asterisks: Student’s *t* tests, *p* = 0.037 and < 0.001, respectively.

We next quantified the abundance of VGLUTs in the vestibular ganglion neurons, which were identified by the expression of neurofilament H and beta-III-tubulin (Nf-H/TuJ1). Counting the spots of both RNAscope probes in the single cryosections through vestibular neurons of *Vglut3*^+/+^ and *Vglut3*^-/-^ mice revealed significantly lower numbers in the *Vglut3*^-/-^ mice for both, *Vglut2* and *Vglut1* transcripts (**Fig. 2I-J**). Since we observed two populations of vestibular neurons, Nf-H/Tuj1-“rich” and “poor” neurons, we performed further analysis subdividing the two populations. Again, a strong reduction in the *Vglut2* signal was observed in the KO vs WT vestibular neurons of NF-H/Tuj1-“poor” and a tendency in NF-H/Tuj1-“rich” population (**Suppl. Fig. 2B-C**), whereas the difference for *Vglut1* was no longer statistically significant upon division of the neurons in two categories. Together, the data suggests that alteration in excitation of the vestibular neurons (due to loss of quantal transmission) may lead to a reduction in neuronal expression of VGLUT1/2 transporters.

As shown above, *Vglut1* and *2* transcripts are robustly expressed in vestibular ganglion neurons, and *Vglut3* transcripts are robustly expressed in vestibular hair cells (McInturff *et al*, 2018; see also **Suppl. Fig. 2A**; data publicly available at umgear.org). Next, we sought to confirm this with confocal immunofluorescence detection of the VGLUT protein isoforms in the same type of tissue sections used for RNAscope. VGLUT1-3 exhibited regional variation in the utricle and crista of *Vglut3*^+/+^ and *Vglut3*^-/-^ (**Fig. 3**). At low magnification, VGLUT3 appeared to be located basally within the sensory epithelia of the utricle and crista. VGLUT1 and 2 immunofluorescence extended from basal locations to the apical surface of the epithelia. Laterally, VGLUT1 tended to be located in the center of the utricle and crista, while VGLUT2 was relatively absent from the center and concentrated in peripheral zones (**Fig. 3A**). The same patterns were observed for VGLUT1 and 2 in *Vglut3^-/-^* (**Fig. 3C**). At high magnification, VGLUT3 was seen localized mainly to the basolateral “tails” of type II VHCs below the nuclei, as well as in type I VHCs (**Fig. 3B**). Based on overlap with the neuronally expressed Na^+^/K^+^ ATPase, VGLUT1 and 2 proteins appeared to be located sparsely in peripheral axons and robustly in peripheral terminals, including the calyces that envelop type I VHCs (**Suppl. Fig. 3**). In addition, VGLUT2 was clearly detected in some type II VHCs in both *Vglut3^+/+^*and *Vglut3^-/-^* (**Fig. 3B, D, Fig. 4**). Due to the small cytoplasmic volume in type I VHCs and the very tight ensheathment of postsynaptic calyces which often express VGLUT2, the expression of VGLUT2 in type I VHCs was difficult to resolve with conventional confocal microscopy. Thus, we next quantified this in thinner optical sections using Airyscan deconvolution confocal microscopy.

**Figure 3.**
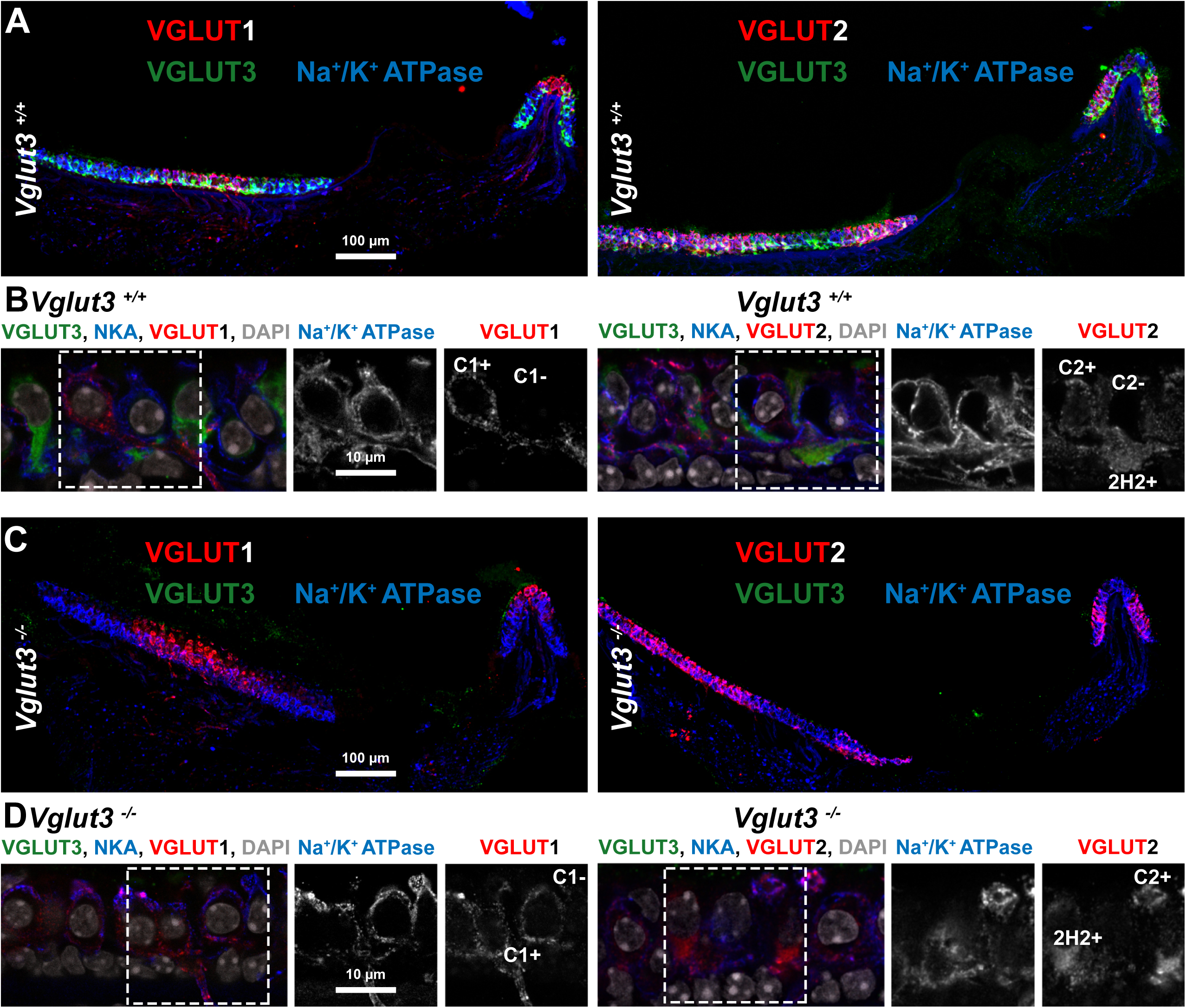
VGLUT1, 2, and 3 protein expression in the utricle and crista. **A)** Cryosections of the utricle and crista from 8-week old *Vglut3^+/+^* mice were labeled with antibodies for VGLUT3, Na^+^/K^+^ ATPase (NKA), and either VGLUT1 (*left*) or VGLUT2 (*right*). **B)** High magnification confocal images of samples as in panel A, plus DAPI stain, for *Vglut3^+/+^* showing VGLUT1 (*left*) or VGLUT2 (*right*). **C)** As in panel A but for *Vglut3^-/-^*. **D)** As in panel B but for *Vglut3^-/-^*. On the right side, grey scale panels show inserts with Na^+^/K^+^ ATPase and VGLUT1 (*left*) or VGLUT2 (*right*). C1+: calyx with VGLUT1; C1-: calyx without VGLUT1; C2+: calyx with VGLUT2; C2-: calyx without VGLUT2; 2H2+: Type II hair cell with VGLUT2.

**Figure 4.**
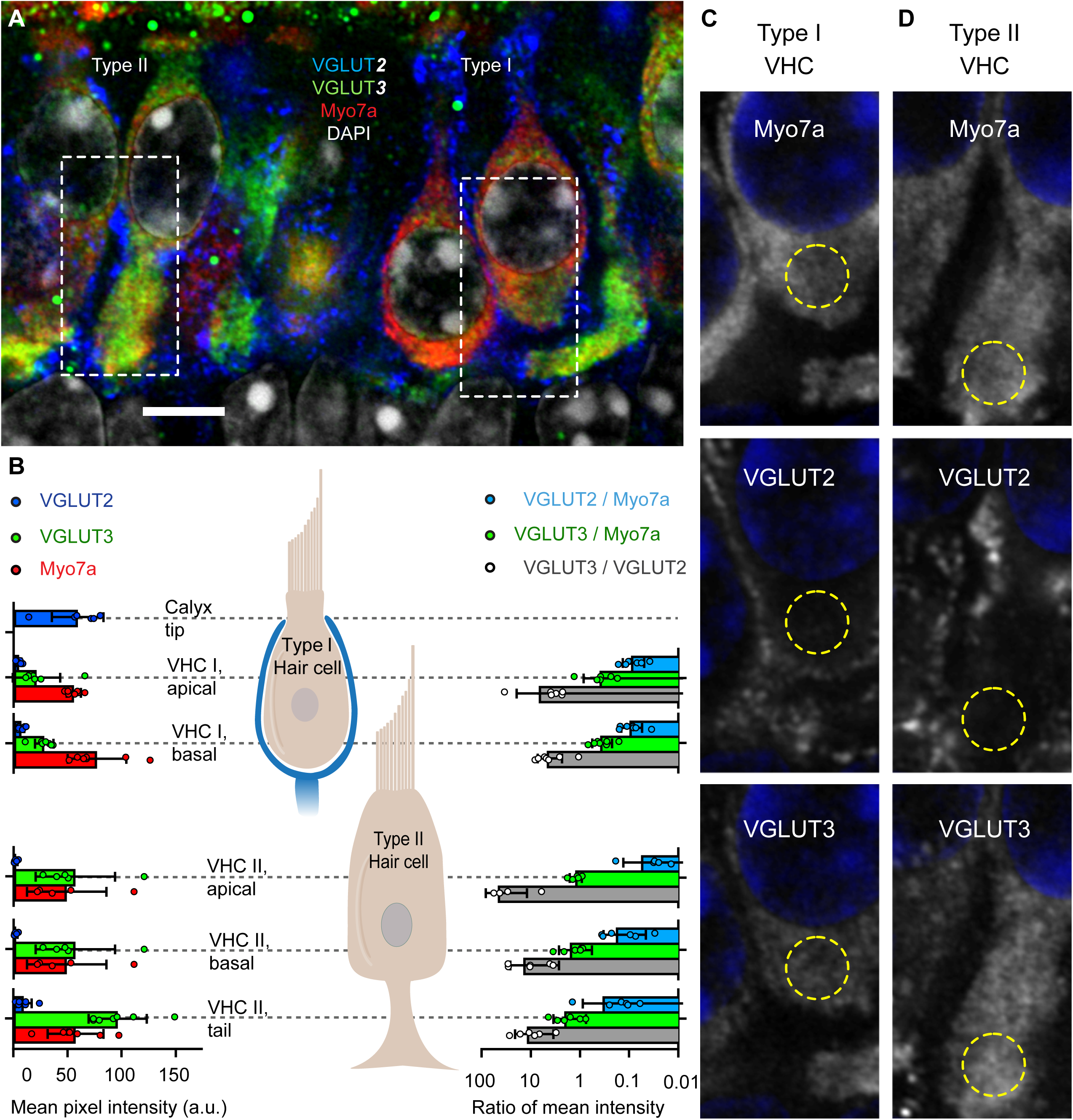
Protein expression profile of VGLUT2 and VGLUT3 in the *Vglut3^+/+^* mouse utricle**. A)** Single optical section of Airyscan microscopy on an exemplary utricular cryosection in the central zone from an 8-week old mouse labelled with DAPI, Myosin7a (VHC marker), VGLUT2, and VGLUT3. Type I VHCs can be well distinguished from the type II cells by the narrow necks, lower position of the nuclei and the absence of the cell foot/tail. Dashed rectangles outline the magnified regions displayed in panels C-D. **B)** Quantification of fluorescence intensity in six regions of interest (*left*). *Right*: Ratios of mean intensity (log scale) for the same 5 hair cell regions. Bars are group means ± SD and points are image means from individual regions of interest; N = 3 mice, 2 – 3 images per utricle. **C)** Magnified regions of the type I hair cell, from top to bottom: Myosin7a, VGLUT2, or VGLUT3 are shown in gray scale; DAPI (*blue*). Yellow circle shows region of interest for the type I VHC below nucleus. **D)** Same as panel C for the type II VHC. Yellow circles show region of interest for the type II VHC tail.

### VGLUT3 is the predominant isoform expressed prominently in vestibular hair cells, with a minor expression of VGLUT2

Immunofluorescence of VGLUT2 and 3 relative to Myosin7a in VHCs from cryosections of the central zone of utricles from 8-week-old mice revealed strong expression of VGLUT3 in type II VHCs, whereas type I VHCs were significantly less immunoreactive (**Fig. 4, Suppl. Fig. 4**). In both, VGLUT3 fluorescence showed similar intensity below and above the nuclei and peak fluorescence was typically observed in the tails of type II VHCs (**Fig. 4B-D, Suppl. Fig. 4**). VGLUT2 fluorescence was consistently much less than VGLUT3, on average, and similarly low across all of the regions of interest in both types of VHCs. The VGLUT3/VGLUT2 fluorescence ratio on average amounted to 6 in type I and 25 in type II VHCs. On the other hand, VGLUT2 fluorescence was robust in the tips of calyces, similar to immunofluorescent levels of VGLUT3 in type II VHCs (**Fig. 4B, Suppl. Fig. 4**). The Myosin7a fluorescence was similar across VHC regions of interest. Relative to Myosin7a, VGLUT2 fluorescence was consistently low, amounting to 10-20%, on average, while VGLUT3 fluorescence was on average around 40% in type I and up to twice the Myosin7a fluorescent levels in type II VHCs, respectively (**Fig. 4B**). Taken together, our RNA and protein expression studies suggest that VGLUT3 is the predominant, if not the only essential vesicular glutamate transporter in VHCs. This is consistent with previous results using single-cell RNA sequencing, which found robust expression of *Vglut3* mRNA in type I and type II VHCs, that the great majority of VHCs do not generate *Vglut2* mRNA at detectable levels, and that *Vglut1* mRNA was absent (McInturff *et al*, 2018; see also **Suppl. Fig. 2A**).

### Zonal variation of VGLUT3 expression in utricular hair cells

When looking throughout the utricle, we noticed large heterogeneity in the VGLUT3 immunofluorescence signal in type I VHCs. To assess the relative expression levels of VGLUT3 in the different zones and VHC types in the mouse utricle, we quantified its expression in the striola and extrastriola. Utricles from *Vglut3^+/+^*mice between the ages of P15-30 (N = 4) were labelled with VGLUT3, calretinin, and beta-III-tubulin. With the help of these zonal markers, we were able to faithfully quantify the expression levels of VGLUT3 in the type I and type II VHCs in the two utricular zones (**Fig. 5A**). We found several populations of type I VHCs - calyx combinations in the striola. Cells surrounded by calretinin and beta-III-tubulin (TuJ1) in the striola were most likely type I VHCs surrounded by calyx-only fibers. Cells that were not surrounded by calretinin but were positive for beta-III-tubulin were likely type I VHCs surrounded by the dimorphic fibers in the striola (Leonard & Kevetter, 2002; Desai *et al*, 2005a, 2005b; Eatock *et al*, 2008). We also observed type I VHCs that were mildly positive for calretinin. Type II VHCs in the extra-striolar region were mildly positive for calretinin, as previously described (e.g. Desai *et al*, 2005a; Eatock & Songer, 2011; Prins *et al*, 2020). VGLUT3 showed variable expression in all these VHCs. We found a few type I VHCs, often surrounded by calretinin-positive calyces that had very weak/almost undetectable VGLUT3 expression. The extrastriolar type I VHCs almost always expressed detectable levels of the transporter. In contrast, type II VHCs had much stronger expression, roughly 10 times stronger than the type I VHCs, in the striola and approximately 2-5 times stronger in the extrastriola (**Fig. 5A-B**). These data collectively suggest that while synaptic transmission in type II VHCs throughout the organ and type I VHCs in the extrastriola is supported by VGLUT3, striolar type I VHCs may not heavily depend on glutamatergic transmission. While we cannot exclude the possibility of low VGLUT3 expression below the threshold of detection, it is questionable whether such scarce expression is sufficient to support glutamatergic transmission at the striolar type I VHCs.

**Figure 5.**
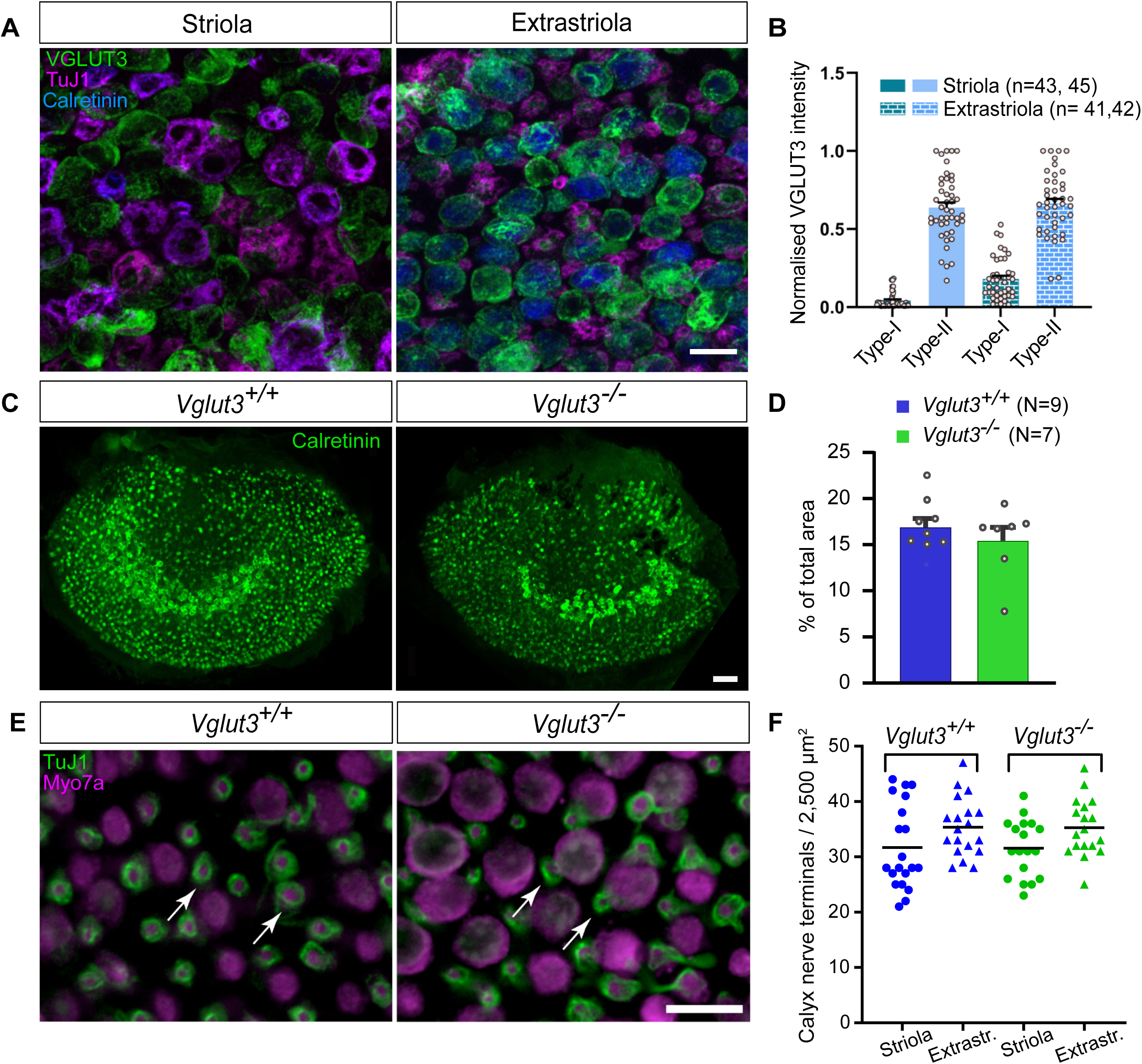
VGLUT3 and Calretinin expression in utricles of *Vglut3^+/+^* and *Vglut3^-/-^*. **A)** Representative images of the VGLUT3 immunosignal (*green*) in the striola and extrastriola of *Vglut3^+/+^* utricles. Utricles were co-labeled with calretinin (*blue*) to distinguish the central and peripheral zones, striola and extrastriola, respectively, as calretinin strongly labels calyceal (but not dimorphic) endings in the striola. Neuronal marker TuJ1 (*magenta*) was used to distinguish between VHC types based on the postsynaptic connections. Scale bars: 10 μm. **B)** Intensity of the VGLUT3 immunosignal normalized to the brightest VHC per image. Type I VHCs show significantly less VGLUT3 immunosignal compared to the type II VHCs, which is further reduced in the striola compared to extrastriola, where the VHC-Is typically seem devoid of VGLUT3. **C)** Representative 20X confocal images displaying the striolar calyces and extrastriolar type II VHCs immunolabeled with calretinin in the *Vglut3^+/+^* and *Vglut3^-/-^* utricles. Scale bars: 50 μm. **D)** Analysis of the calretinin-positive striolar area normalized to the area of the entire organ in the *Vglut3^+/+^* and *Vglut3^-/-^* utricles of young mice (P15-30), showing no obvious differences in the striolar width/size in these two genotypes. **E-F)** Counting the number of calyces in two 50 x 50 µm regions in the striolar and medial extrastriolar regions revealed no difference in calyx numbers among the two genotypes. Data consist of 20 samples / region from 10 utricles (4-12 months of age). Scale bar: 10 µm. Arrows point at exemplary calyces.

Calretinin is known to strongly label a vast majority of striolar calyces (e.g. Desai *et al*, 2005a; Eatock & Songer, 2011; Leonard & Kevetter, 2002; Hoffman *et al*, 2018; Prins *et al*, 2020). To unveil possible effects of VGLUT3 deletion on preservation of striolar calyces, we additionally inspected the calretinin immunofluorescence signal in the striolar area of the utricles. We observed no systematic difference in the striolar width positive for calretinin in the utriculi of *Vglut3^-/-^* mice and the respective wild-type controls (**Fig. 5C-D**), suggesting no overt loss of striolar calyceal afferents, consistent with the observations of Regalado Núñez and colleagues (2024). Unlike observations in the cochlea, where all inner hair cells robustly express VGLUT3 to a similar degree and loss of VGLUT3 leads to synapse loss and neuronal degeneration already in young mice (Seal *et al*, 2008; Ruel *et al*, 2008; Kim *et al*, 2019), even at 4-12 months of age, we observed similar utricular calyx numbers between the two genotypes (**Fig. 5E-F**). Since ca 85% of vestibular afferents possess calyx terminals (either calyx-only or dimorphic afferents; Kevetter & Leonard, 2002; Lysakowski *et al*, 1999; Fernandez *et al*, 1988, 1990), our results suggest that (at least) 80-85% of vestibular afferents survive in *Vglut3^-/-^*mice.

### The absence of VGLUT3 results in > 95% reduction in EPSC rate between type I VHCs and calyces

To measure quantal release from the type I hair cell-calyceal synapses in the utricle we recorded excitatory post-synaptic currents (EPSCs) from the complex calyces and a few simple calyces in the extrastriolar and striolar regions. The mean capacitance of the calyces recorded was 15.4 ± 2.0 pF, ranging from roughly 6 pF for a simple calyx to approximately 43 pF for a complex dimorphic calyx. The average series resistance recorded was 32.0 ± 2.9 MΩ. Calyces were first held at −79 mV and spontaneous EPSCs were recorded with 5.8 mM K^+^ in the extracellular solution (**Fig. 6**). Under these conditions, 25% of the *Vglut3^+/+^* calyces were silent and did not show any spontaneous EPSCs and 75% showed very few. The mean frequency under these conditions was 0.10 ± 0.02 Hz. Hair cells were depolarized with 22 mM K^+^ in the extracellular solution, which increased the EPSC frequency to 10.95 ± 2.21 Hz in *Vglut3^+/+^* calyces (**Fig. 6B**). Evoked EPSCs in the *Vglut3^+/+^*calyces were divided into two broad categories of simple and complex events. We further divided the simple events into single or double exponential events depending on their decay kinetics (**Fig. 6C**). Single exponential decay time constants ranged from 0.25 to 10.70 ms. Double exponential events had a fast decay followed by a slower one, where the time constants were separated by, at least, a factor of 10. The average time constants for the fast (range: 0.13 to 2.65 ms) and slow components (range: 2.05 to 32.96 ms) of the events with two decays were 0.63 ± 0.02 ms and 9.05 ± 0.44 ms, respectively. In contrast, the calyces of *Vglut3^-/-^* mice showed almost no EPSCs in 5.8 mM (0.004 ± 0.004 Hz) or 22 mM K^+^ (0.39 ± 0.33 Hz) (**Fig. 6D-F**). We note that these events had small amplitudes, comparable to the recording noise and did not follow the typical synaptic current waveform of a fast rise and an exponential decay.

**Figure 6.**
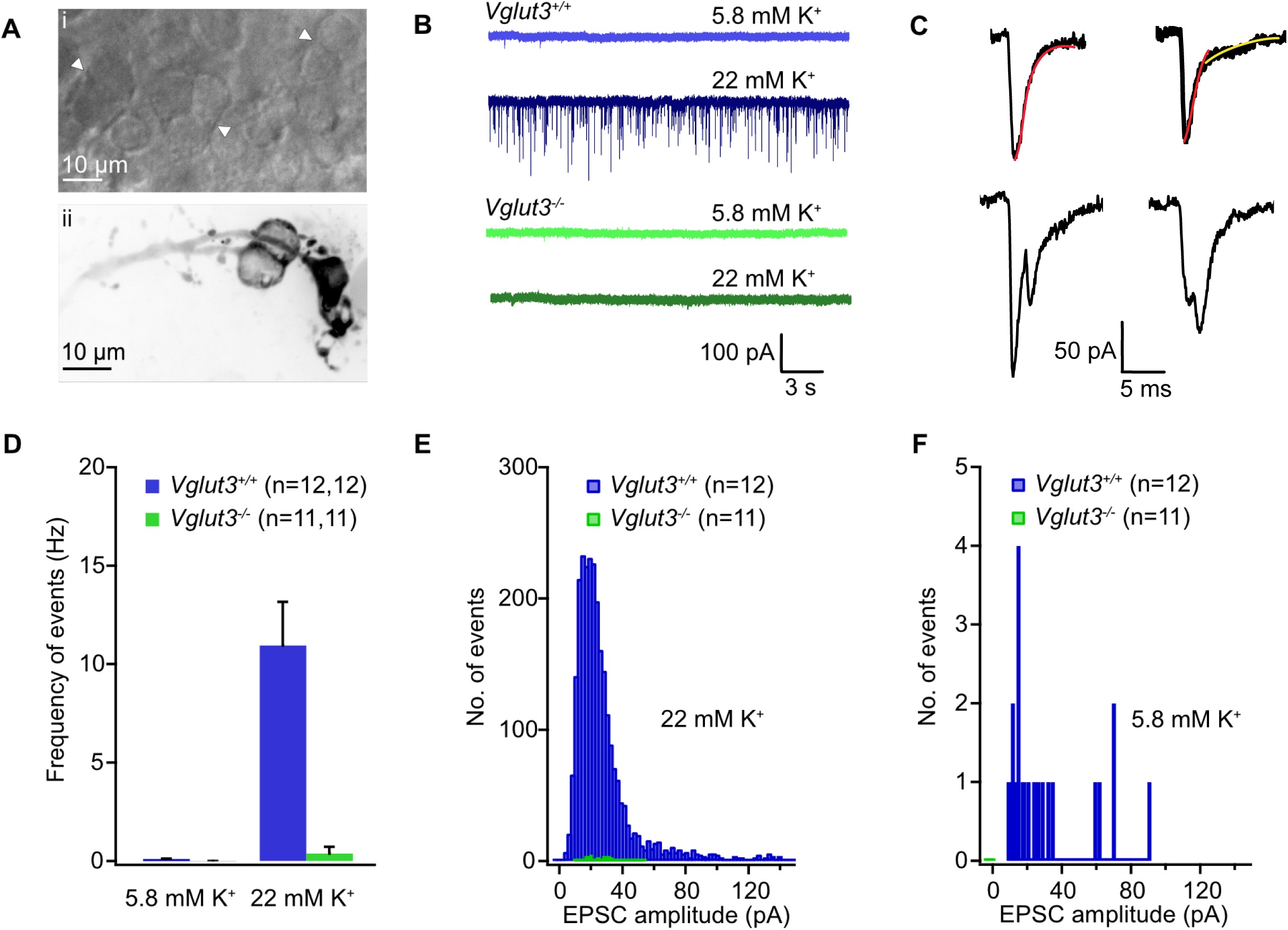
EPSC loss in the calyces of *Vglut3^-/-^* utriculi. **Ai)** A bright-field image of the striolar region of a utricle at 60x magnification showing complex calyces (white arrowheads). **Aii)** A complex dimorphic calyx filled with Alexa 568 hydrazide dye showing several boutons and two double calyces originating from a single fiber. **B)** Example postsynaptic currents recorded from calyces held at −79 mV in the presence of low or high extracellular [K^+^]. Numerous EPSCs were elicited in response to 22 mM K^+^ in the *Vglut3^+/+^* calyces as opposed to *Vglut3^-/-^* calyces. **C)** Events were broadly classified into simple single exponential (*left*), or simple double exponential (*right*), or complex events (*bottom*). **D)** Histogram of the frequency of events generated in response to 5.8 mM K^+^ and 22 mM K^+^ in the *Vglut3^+/+^* and *Vglut3^-/-^*calyces shows a very strong reduction (approx. 10 times) of events in the calyces lacking presynaptic VGLUT3. **E-F)** EPSC amplitude distributions for the events recorded in evoked (22 mM K^+^) and resting (5.8 mM K^+^) conditions showing most *Vglut3^+/+^*amplitudes ranged from 15 to 40 pA, whereas VHCs in *Vglut3^-/-^*failed to elicit enough events in both conditions.

Similar results were obtained using a holding potential of −103 mV and depolarization with 40 mM K^+^, to elicit larger and more frequent EPSCs (**Fig. 7**). Under these conditions, roughly 8% of *Vglut3^+/+^* calyces were silent, and elicited events with a mean frequency of 0.53 ± 0.20 Hz at 5.8 mM K^+^. Stimulation with 40 mM K^+^ led to a significant increase in the mean frequency to 14.94 ± 3.92 Hz, whereas EPSC rates remained very low in *Vglut3^-/-^* calyces when VHCs were depolarized in either 5.8 mM (0.07 ± 0.03 Hz) or 40 mM K^+^ (0.43 ± 0.25 Hz). In both evoked conditions (22 or 40 mM K^+^) we detected a large number of events with slower decay rates in *Vglut3^+/+^*, which is directly correlated to glutamate spillover and its rate of clearance from the narrow synaptic cleft (Sadeghi *et al*, 2014). The average amplitudes of events generated were 30.75 ± 0.50 pA and 55.34 ± 0.63 pA in the 22 mM and 40 mM [K^+^] conditions, respectively.

**Figure 7.**
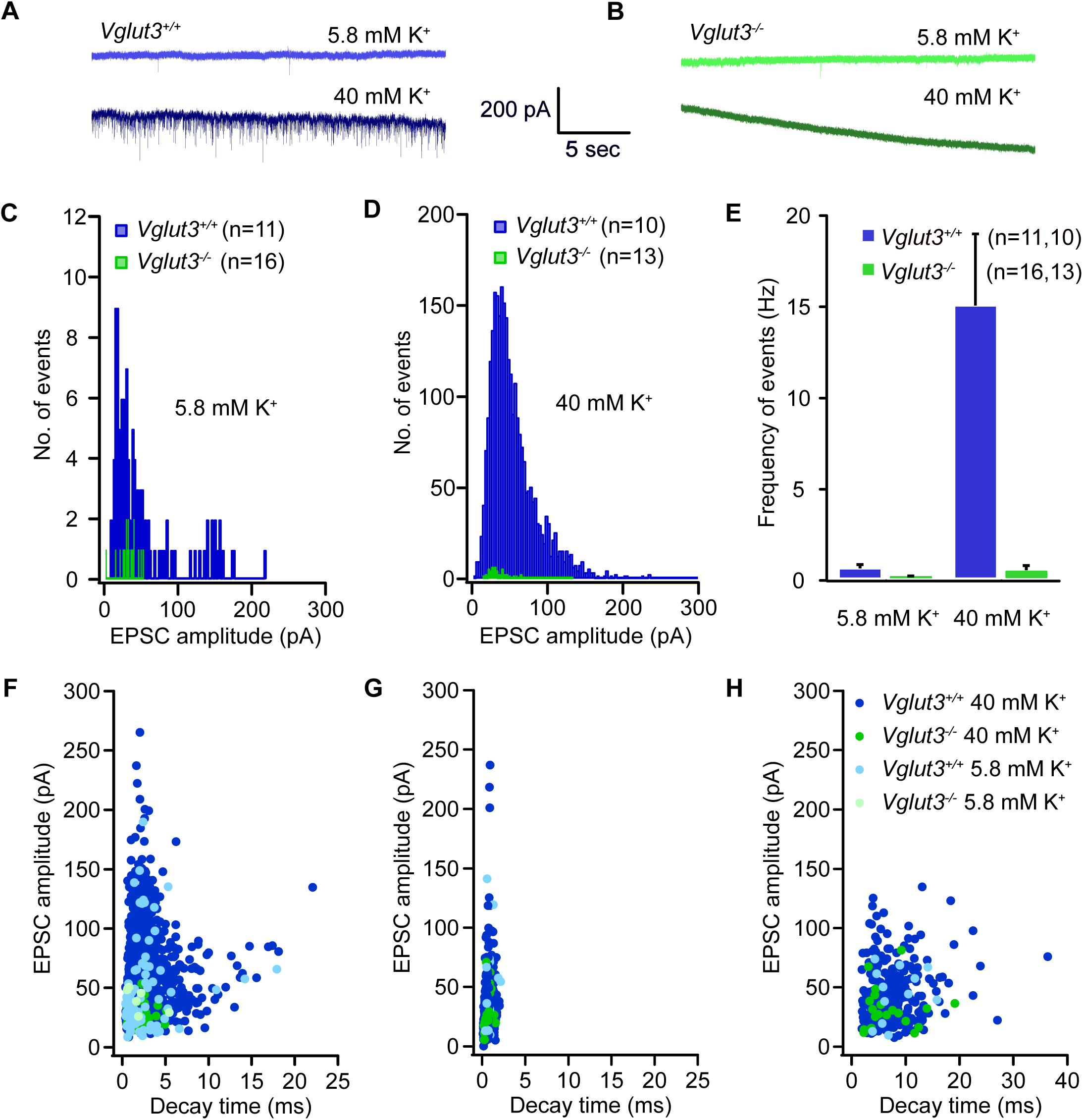
No increase in postsynaptic activity upon stronger VHC activation. **A-B)** Example postsynaptic current recorded in the resting (5.8 mM K^+^) and evoked (40 mM K^+^) conditions in the calyces of *Vglut3^+/+^* (A) and *Vglut3^-/-^* (B) mice at −103 mV holding potential. **C-D)** Amplitude distribution of postsynaptic events at 5.8 mM K^+^ (C) and 40 mM K^+^ (D) showing the larger portion of events in both conditions ranged between 15-100 pA amplitude in the *Vglut3^+/+^* calyces, while the *Vglut3^-/-^*calyces had limited events with smaller amplitudes. **E)** Histogram with EPSC frequency in both conditions and genotypes displays the failure to elicit events in the *Vglut3^-/-^* calyces even upon stimulation by 40 mM K^+^. **F-H)** Scatter plots of EPSC amplitudes with their respective decay times for the simple events with single exponential decays (F), and the fast (G) and slow (H) component of events with double exponential decays, where the slow and fast components were separated by at least a factor of 10.

Together, the data suggest that glutamatergic signaling in calyces depends on the presence of VGLUT3. The lack of convincing EPSCs in *Vglut3^-/-^*suggests that these utricular calyces completely lack glutamatergic quantal synaptic transmission.

### Evoked action potentials in calyces ex vivo and spontaneous action potentials in vestibular afferent neurons in vivo are similar in *Vglut3^+/+^* and *Vglut3^-/-^*

To study the effects of VGLUT3 deletion from hair cells on the excitability of postsynaptic calyx-bearing vestibular neurons, we studied action potential generation in response to step current injection in the *Vglut3^+/+^*and *Vglut3^-/-^* utricles. Current steps of +10 pA were injected into the calyces to evoke spikes. Most calyces in either genotype did not show any spontaneous spiking in our recording conditions. Evoked spikes were categorized into two classes based on earlier reports (Curthoys *et al*, 2017; Kalluri *et al*, 2010; Eatock & Songer, 2011; Goldberg & Fernandez, 1971). Sustained spiking calyces generated action potentials throughout the current step, whereas transient calyces spiked once or twice at the onset of the current injection, and occasionally at the end of the current step as a rebound spike. **Figure 8A-C** shows representative traces of sustained and transiently spiking calyces in the *Vglut3^+/+^*and *Vglut3^-/-^* utricles. All the calyces in this dataset belonged to dimorphic fibers originating from the striola or extra-striolar region. In 8 of 10 calyces in the *Vglut3^+/+^* utricles and in 8 of 14 calyces in *Vglut3^-/-^* we recorded sustained firing. In the remaining calyces transient firing was detected, indicating they belonged to striolar dimorphic nerve fibers. It has been reported earlier that transient neurons have higher current thresholds at rheobase, more negative resting potentials, and lower input resistances than sustained units (Kalluri *et al*, 2010; Risner & Holt, 2006) due to their large resting K^+^ conductances. Comparing the sustained units obtained from each genotype (**Fig. 8D-I**), the mean resting potential was not significantly different between *Vglut3^+/+^*(−70.2 ± 1.3 mV) and *Vglut3^-/-^* (−71.5 ± 2.6 mV). Other indicators of intrinsic excitability of a cell like rheobase and voltage threshold also did not differ significantly between the two groups with threshold values of −57.1 ± 2.4 mV and −60.3 ± 1.1 mV for the *Vglut3^+/+^*and *Vglut3^-/-^*, respectively. First spike latency at rheobase: *Vglut3^+/+^* (167 ± 50 ms); *Vglut3^-/-^* (188 ± 75 ms) and maximum evoked spike rate: *Vglut3^+/+^* (42.3 ± 7.5 Hz); *Vglut3^-/-^* (50.9 ± 10.6 Hz) were similar, as well.

**Figure 8.**
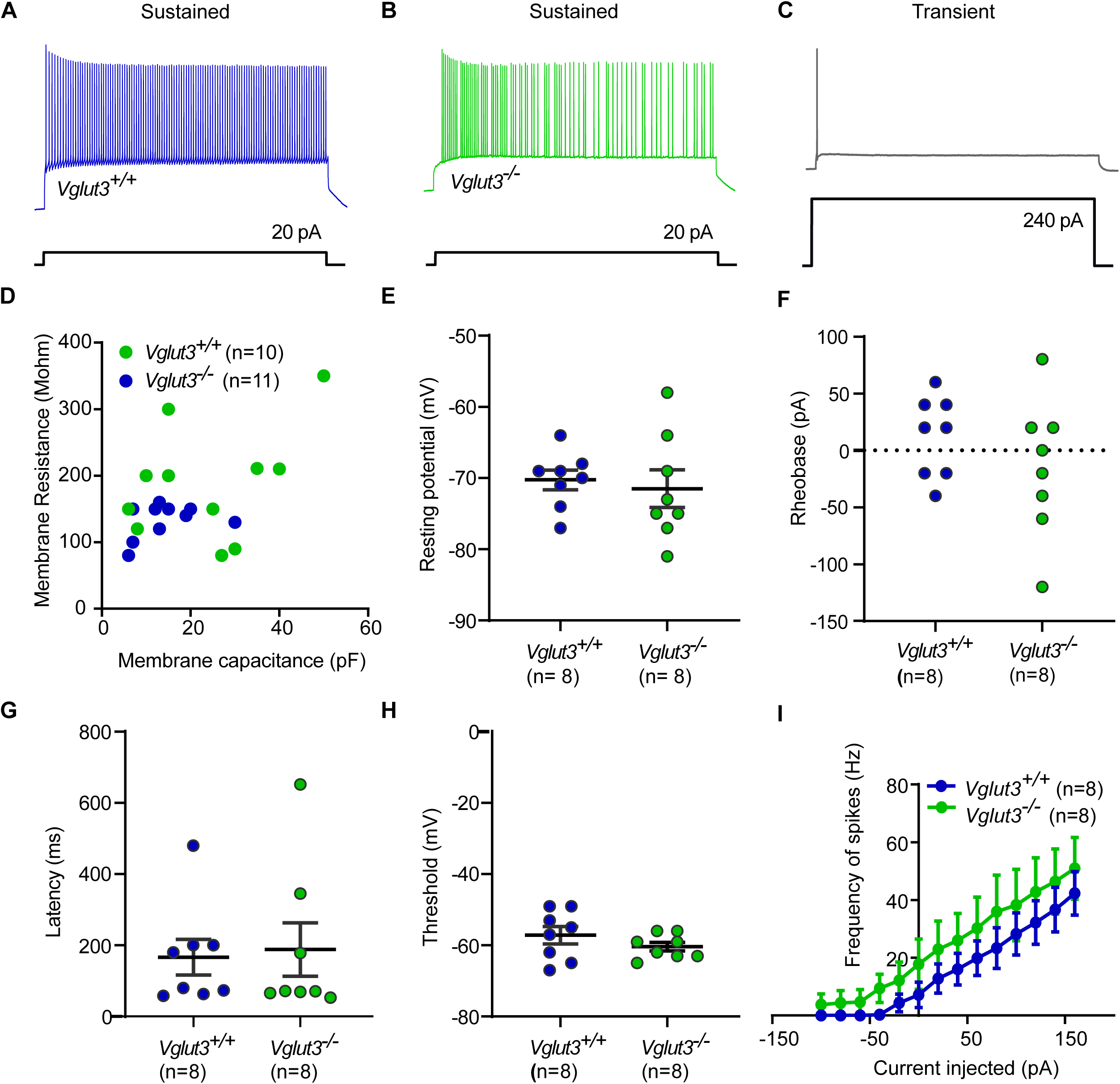
Action potentials analysis in the vestibular calyces of *Vglut3^+/+^* and *Vglut3^-/-^* mice. **A-C)** Representative traces of sustained units from the *Vglut3^+/+^* and *Vglut3^-/-^*calyces and one *Vglut3^+/+^* transient unit (C). **D)** Scatter plot of the membrane resistance and capacitance for the two genotypes **E)** Resting potentials measured at current clamp with I= 0. **F-H)** Histograms of intrinsic excitability parameters of the sustained units such as (F) rheobase/current threshold, (G) latency of the first action potential, and (H) voltage threshold to generate action potential, showed no significant differences between the *Vglut3^+/+^* and *Vglut3^-/-^*calyces. **I)** Frequencies of spikes versus the currents injected were not significantly different between the sustained *Vglut3^+/+^* and *Vglut3^-/-^* calyces, although there was a trend of marginally higher frequencies for the *Vglut3^-/-^* units.

To ask if the absence of quantal transmission from VHCs eliminated or altered spontaneous activity of the vestibular nerve, we next recorded spike rates and the regularity of inter-spike intervals *in vivo*. Example spike trains and inter-spike interval histograms are shown in **Suppl. Figure 5**. These single unit recordings revealed apparently normal spontaneous spiking with no obvious differences in the distributions of spike rates, inter-spike-intervals, or coefficient of variation (CV) of inter-spike-intervals (**Fig. 9**). Group means of mean spike rates were 63.8 ± 48.1 Hz and 40.0 ± 25.3 Hz for *Vglut3^+/+^* and *Vglut3^-/-^*, respectively (*p* = 0.90; Wilcoxon rank sum test). The mean CVs were 0.41 ± 0.43 and 0.42 ± 0.36, respectively (*p* = 0.53). Our data collectively show unaltered excitability of afferent vestibular neurons, implying it is running independent of quantal transmission.

**Figure 9.**
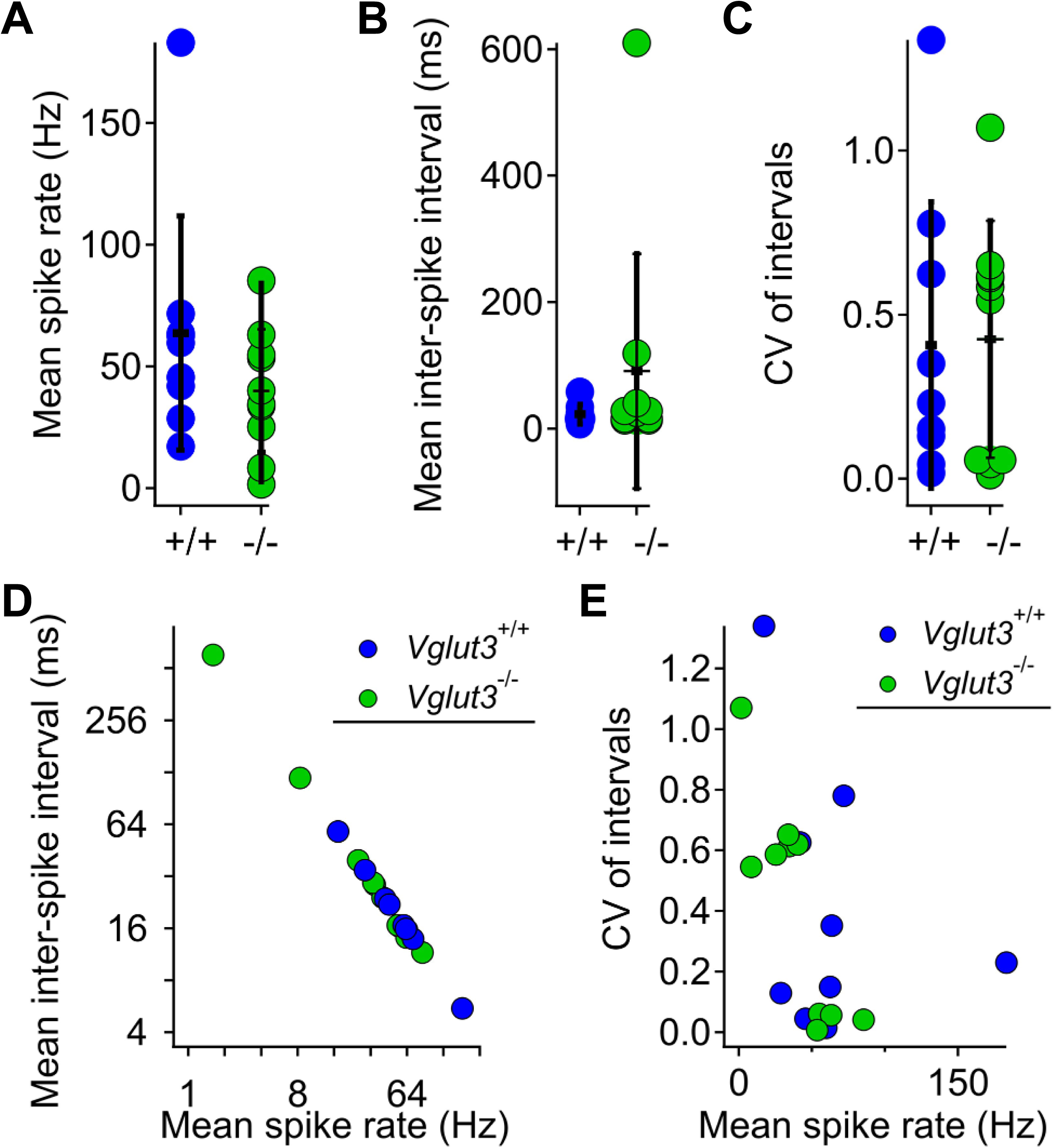
Apparently normal spontaneous activity in the vestibular neurons of *Vglut3^-/-^* mice. **A-C)** Distributions of mean spike rate (A), inter-spike-interval (B), and CV of intervals (C). Horizontal lines are mean, vertical lines are ± SD. N = 9 *Vglut3^+/+^* and 10 *Vglut3^-/-^* units. **D)** Mean inter-spike-interval vs mean spike rate for 9 WT and 10 KO units. **E)** CV of intervals vs mean spike rate.

To illustrate the contrast between apparently normal vestibular nerve spontaneous activity and our expectation of completely absent auditory nerve spontaneous activity in *Vglut3^-/-^*, we recorded from auditory nerve fibers (ANF) with and without an applied sound stimulus (**Fig. 10**). Each ANF showed spontaneous activity in recordings of 28 units from 10 *Vglut3^+/+^*mice (mean 32.3 ± 34.0; range 1.5 to 120.5 Hz). In contrast, in 10 *Vglut3^-/-^*mice with a similar number of recording attempts, we encountered only 3 units with spontaneous activity (mean spike rates of 11.6, 17.2, and 55.5 Hz; **Fig. 10A-B**). Compared with *Vglut3^+/+^* ANF units of similar mean rate, inspection of cumulative inter-spike interval histograms of the three “responsive” units from *Vglut3^-/-^* mice revealed a less sigmoidal shape due to absence of short inter-spike intervals (**Fig. 10C**). The characteristic frequency (CF) of each *Vglut3^+/+^* ANF was identified, and CF tone bursts were presented at different levels. These typically evoked monotonically increasing spike rates with increasing sound pressure level (SPL). Accompanying peri-stimulus time histograms showed the classic pattern of rapid spike onsets, followed by slower adaptation. In contrast, spike rates of *Vglut3^-/-^*ANFs were not modulated by noise bursts at any SPL (**Fig. 10D-H**). We compared the CVs of inter-spike intervals (ISIs) of evoked spike trains in *Vglut3^+/+^* ANFs and cochlear nucleus units with the CVs of ISIs of spontaneous spike trains assessed in Fig. 10C for the three presumed ANFs in *Vglut3^-/-^* and three *Vglut3^+/+^* ANFs of similar spontaneous rate (**Fig. 10I**). Spontaneous spike trains in both *Vglut3^+/+^* and *Vglut3^-/-^* were more variable than evoked spike trains in *Vglut3^+/+^*, and 2 of the 3 spontaneous spike trains from the presumed ANFs in *Vglut3^-/-^*displayed the greatest variability.

**Figure 10.**
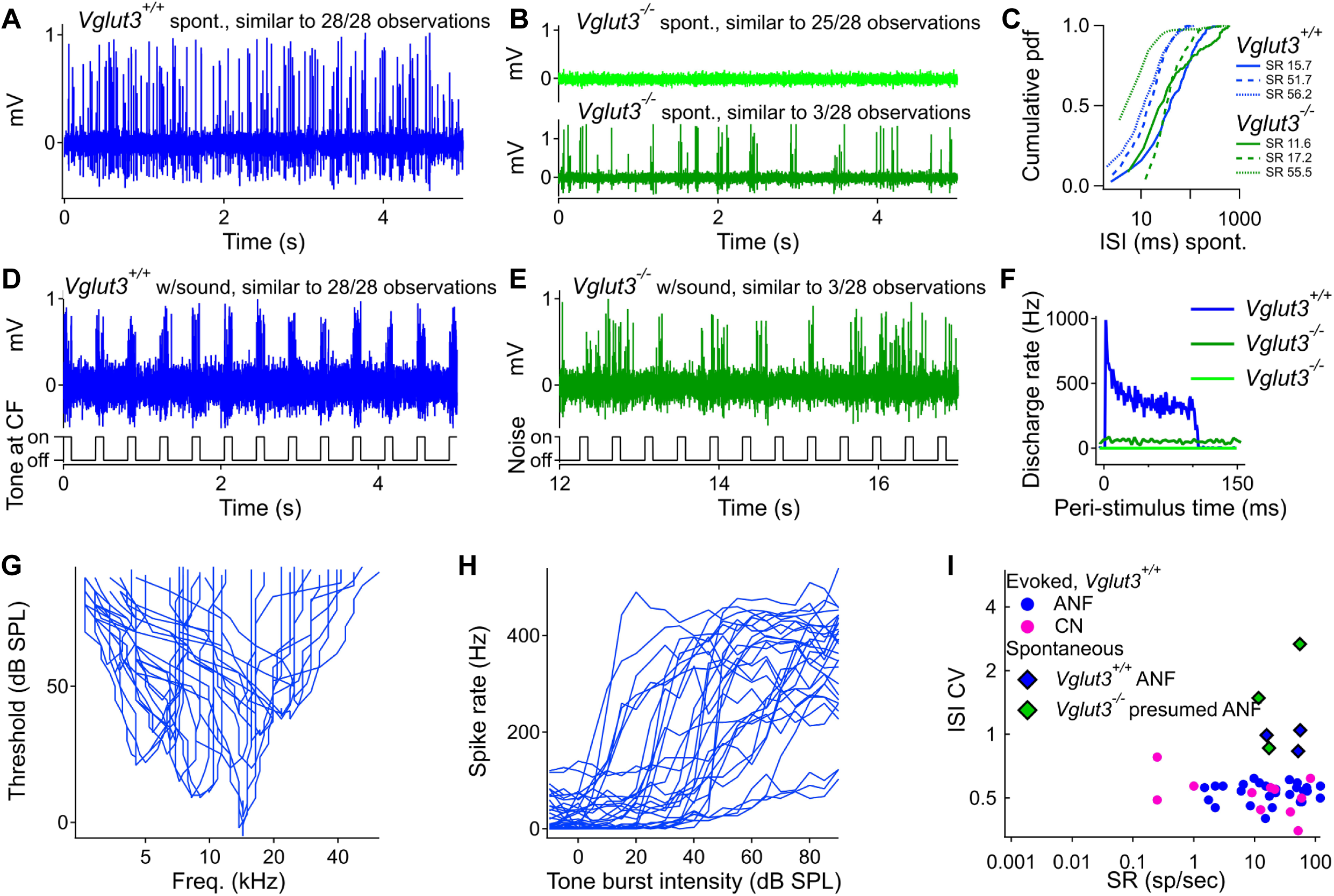
Single unit recordings of auditory nerve fibers in *Vglut3^+/+^* and *Vglut3^-/-^* mice. **A)** In *Vglut3^+/+^* mice, spontaneous activity was observed in all units, showing one example of 28 recordings. **B)** In *Vglut3^-/-^* mice, we typically observed no spontaneous activity. Rarely (3 of 28 recordings), we observed spontaneous activity presumed to originate from type-1 ANFs. **C)** Cumulative probability density function histograms compare the inter-spike intervals for three *Vglut3^+/+^* and *Vglut3^-/-^* units. *Vglut3^-/-^*units generally had lower rates with fewer short intervals, and a greater proportion of long intervals. **D)** In *Vglut3^+/+^* mice, tones at characteristic frequency (CF) evoked spikes that were synchronized to the stimulus in all cases. **E)** In *Vglut3^-/-^* mice, when spontaneous activity was present, spikes were not synchronized to the sound stimulus, which was broadband noise. **F)** Peri-stimulus time histograms for one *Vglut3^+/+^* unit stimulated with 90 dB SPL tones at CF, one *Vglut3^-/-^* unit that was spontaneously active and presented with 90 dB SPL noise at 4-60 kHz, and one *Vglut3^-/-^* unit that was not spontaneously active. **G)** Threshold tuning curves for the 28 *Vglut3^+/+^* recordings. **H)** Rate vs. level functions for the *Vglut3^+/+^* recordings. **I)** A plot of coefficient of variation of inter-spike-intervals (ISI CV) vs spontaneous rate (SR) enabled comparison of activity under four different conditions: evoked spikes in *Vglut3^+/+^* mice (separated into ANF and cochlear nucleus, CN, groups), and spontaneous spikes in *Vglut3^+/+^*and *Vglut3^-/-^* mice. Spontaneous spike timing was more irregular than evoked spikes, in general. Two of the three presumed ANFs in *Vglut3^-/-^*mice (green diamonds, spontaneous activity) exhibited the most irregular spike timing. N = 10 *Vglut3^+/+^* and 10 *Vglut3^-/-^*mice.

### Voltage-gated ion channels in VHCs and calyces of vestibular afferents in the absence of glutamatergic synaptic transmission

The loss of glutamatergic transmission may lead to changes in ion channel expression, so we investigated the expression profile of some prominent channels of the vestibular calyceal synapse. Nonquantal transmission was described to involve primarily the presynaptic low-voltage activated K^+^-conductances, collectively designated as g_K,L_, and postsynaptic K_v_7 and K_v_1 channels (reviewed in Eatock, 2018; Govindaraju *et al*, 2023; Martin *et al*, 2024). We focused our analysis on the KCNQ4 (K_V_7.4) and KCNQ5 (K_V_7.5) channels, expressed in vestibular calyces of adult rodents (Lysakowski *et al*, 2011; Hurley *et al*, 2006; Kharkovets *et al*, 2000; Spitzmaul *et al*, 2013), as well as hyperpolarization-activated cyclic nucleotide-gated (HCN) channels, which influence resting membrane potential and shape postsynaptic action potentials (Horwitz *et al*, 2014, 2011; Rennie & Streeter, 2006; Govindaraju *et al*, 2023; Ventura & Kalluri, 2019).

We immunostained utricles of 2.5- to 5-week-old *Vglut3^+/+^*and *Vglut3^-/-^* mice with KCNQ4, KCNQ5(−2,3), and HCN1 antibodies. According to the manufacturer’s data sheet, the KCNQ5 antibody we used cross reacts with isoforms 1, 2, and 3. Since KCNQ1 (but not KCNQ2 and 3) is reportedly absent from the utricle (Lysakowski *et al*, 2011), we label it as KCNQ5(−2,3). We focused on the type I VHC-calyceal pairs, recognized based on the beta-III-tubulin staining as in **Figure 5B**, and analyzed the mean intensity of these channels in the striola and extrastriola (**Fig. 11A-D**). Striolar analysis included both calretinin– positive and –negative calyces as we observed no systematic differences between these populations. In line with the previous studies (Spitzmaul *et al*, 2013; Hurley *et al*, 2006), KCNQ4 was more strongly expressed in the striola when compared to the extrastriolar region. At the ages tested (between P17 and P35), the origin of the KCNQ4 signal has mostly been ascribed to the inner calyceal membranes rather than type I VHCs (Hurley *et al*, 2006; Spitzmaul *et al*, 2013), which were shown to downregulate the KCNQ4 expression upon maturation whereas developmental upregulation was observed in the calyces. In the *Vglut3^-/-^* utricles, the KCNQ4 intensity appeared lower than controls, however this difference only reached significance in the extrastriola (**Fig. 11D**). Lower immunosignal in the extrastriolar calyces could possibly imply developmental delay in the absence of VGLUT3. KCNQ5(−2,3) expression levels were comparatively low in the striola, and no significant genotype differences were detected. Whereas HCN1 is mostly expressed on hair cell membranes (Horwitz *et al*, 2011), HCN2 is a major calyx channel (Horwitz *et al*, 2014), with HCN1 contributing rather little to the calyceal HCN current (Horwitz *et al*, 2014). HCN1 immunostaining showed relatively low expression levels in the striola than in the extrastriola and was significantly reduced in both regions of *Vglut3^-/-^* relative to *Vglut3^+/+^*. HCN2 immunostaining with different antibodies, however, did not yield satisfactory immunosignals. Therefore, we did not pursue further analysis and turned to electrophysiology.

**Figure 11.**
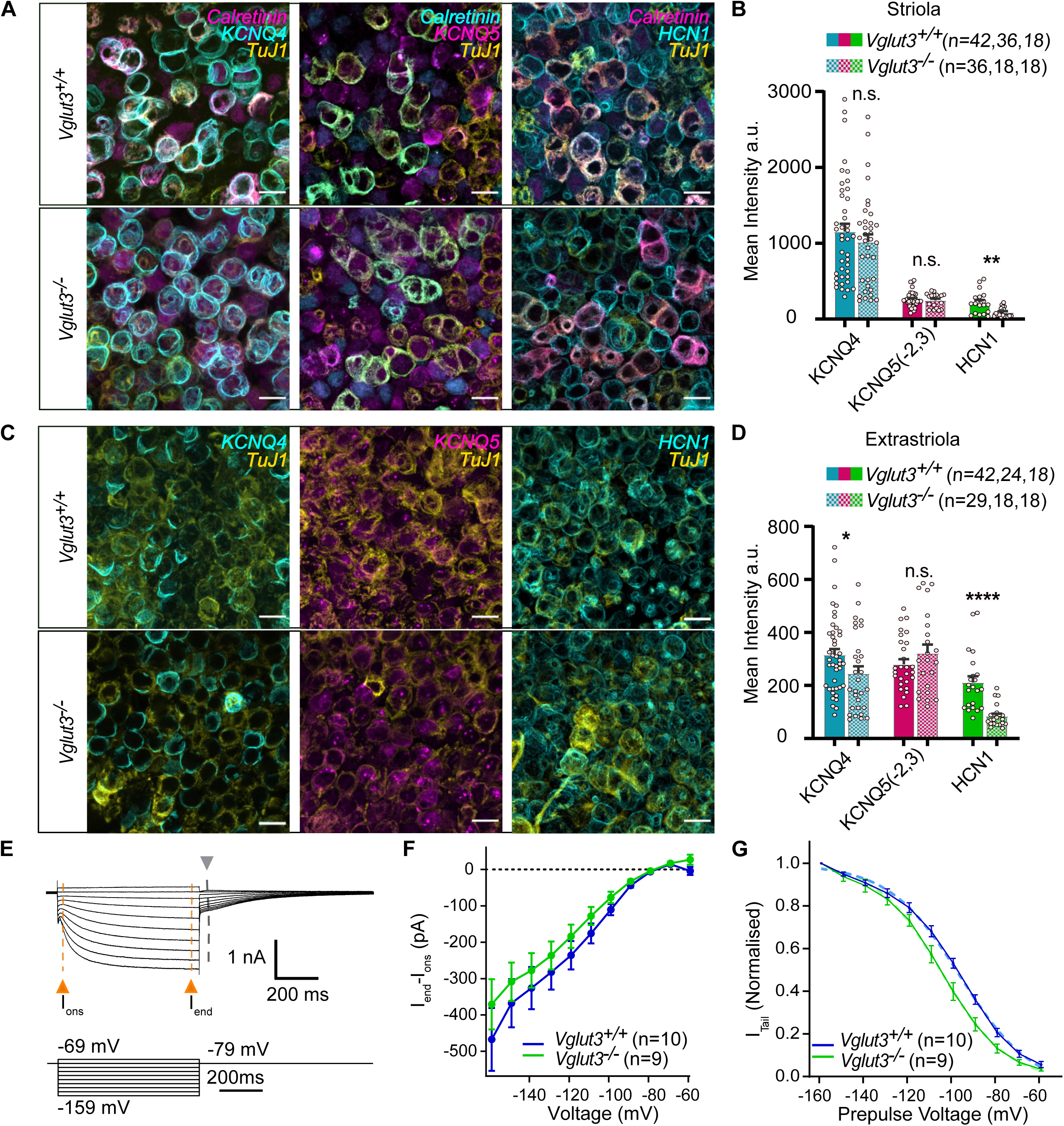
Changes in the expression levels of select ion channels. **A)** Representative immunofluorescence images with calretinin and TuJ1 as context and regional markers and ion channels KCNQ4, KCNQ5 (−2,3), and HCN1 in the *Vglut3^+/+^*and *Vglut3^-/-^* striolas. **B)** Mean intensity analysis for these channels in the striola showing a significant reduction in the HCN1 immunofluorescence in the *Vglut3^-/-^* type I VHCs, while KCNQ4 and KCNQ5 showed negligible differences in the two genotypes. **C)** Same as A, but in the extrastriola, with TuJ1 as a neuronal marker to identify type I VHCs. All scalebars: 10 μm **D)** The mean intensity analysis for the potassium channels in the extrastriola showing a significantly lower intensity of the KCNQ4 and HCN1 immunofluorescence in the calyces and type I VHCs, respectively. **E)** A representative trace showing the HCN2 currents elicited in the calyces in response to hyperpolarizing voltages. The amplitude of the I_end_, I_ons_, and tail currents were extracted from the traces, as indicated in the panel with grey and orange arrows, respectively. **F-G)** The I_end_ – I_ons_ (F) and the normalized tail currents (G) plotted against the prepulse potential. Boltzmann fits of the tail currents reveal significant hyperpolarized shift of ∼ 8-9 mV in V_half_ in the *Vglut3^-/-^* calyces (*p* < 0.0001, extra sum-of-squares *F* test).

We recorded I_h_ with patch-clamp, a current carried mainly through HCN2 channels in the vestibular calyces (Horwitz *et al*, 2014; Meredith *et al*, 2012, 2023; Govindaraju *et al*, 2023). I_h_ or HCN currents are slow activating, non-inactivating currents with a half maximal activation between ca. −130 mV and −100 mV (Horwitz *et al*, 2011, 2014; Meredith *et al*, 2012, 2023; Ventura & Kalluri, 2019; Almanza *et al*, 2012; Bronson & Kalluri, 2023; Chabbert *et al*, 2001), and relative permeability of Na^+^/K^+^ of 1:3-1:5 (Biel *et al*, 2009; Meredith *et al*, 2012). Representative traces showing tail currents are shown in **Figure 11E**. These tail currents were then plotted against the prepulse potential (**Fig. 11G**). By subtracting steady-state current (I_end_) from the onset, peak current (I_ons_; see also Methods and **Fig. 11E**), we further analyzed the total currents (K^+^ and Na^+^) carried by the HCN channels at hyperpolarized potentials (**Fig. 11F**). This also gets rid of any residual contaminating I_KL_ currents, as they have faster activation and inactivation than the HCN currents. While we observed no significant differences in the absolute I_h_ current amplitudes at strongly hyperpolarized voltages (**Fig. 11F**), the tail current analysis revealed a significant shift in the voltage activation of the I_h_ currents in the calyces of *Vglut3^-/-^* animals (**Fig. 11G**). On average, the V_1/2_ of the latter were shifted by −8.64 mV (from −95.86 for *Vglut3^+/+^* to −104.5 in *Vglut3^-/-^*). A hyperpolarized shift in activation could occur due to the loss of HCN1 from the calyces (Horwitz *et al*, 2014) or changes in HCN2 properties. While the more prominent presence of HCN1 in the type I VHCs suggests the observed reduction in the HCN1 immunosignal is most likely related to the loss of HCN1 from the VGLUT3-deficient VHCs, a loss from the VHCs alone or both, calyces and VHCs, is compatible with our electrophysiological data. Recordings made from the striola and extrastriola were pooled, however, 37% of the cells recorded were dimorphic calyces from the striola and 63% belonged to the dimorphic calyces from the extrastriola.

### Modest reduction in synaptic ribbon number in extrastriolar type I VHCs in the absence of glutamatergic transmission

The presynaptic ribbon harbors a halo of synaptic vesicles close to the release sites. To probe whether the loss of glutamatergic transmission affects synaptic ribbon number like in the cochlea, the utriculi of 3-week-old animals of both genotypes were immunostained for the presynaptic ribbon marker Ribeye, the zonal marker calretinin and the neuronal marker beta-III-tubulin or, in some cases, a combination of KCNQ4 and Tenascin. This combination helps distinguish between the VHC types and the utricular zones (**Fig. 12**). We then counted the number of presynaptic ribbons in type I and type II VHCs in both the regions. The quantification revealed a reduction in the abundance of ribbons in type I and type II VHCs of the extrastriola as well as type I VHCs of the striola of *Vglut3^-/-^*animals at P15-39 (**Fig. 12E-F**). This is similar to the effects in the cochlea, where ribbon degeneration is observed upon loss of VGLUT3 or otoferlin (Kim *et al*, 2019; Stalmann *et al*, 2021).

**Figure 12.**
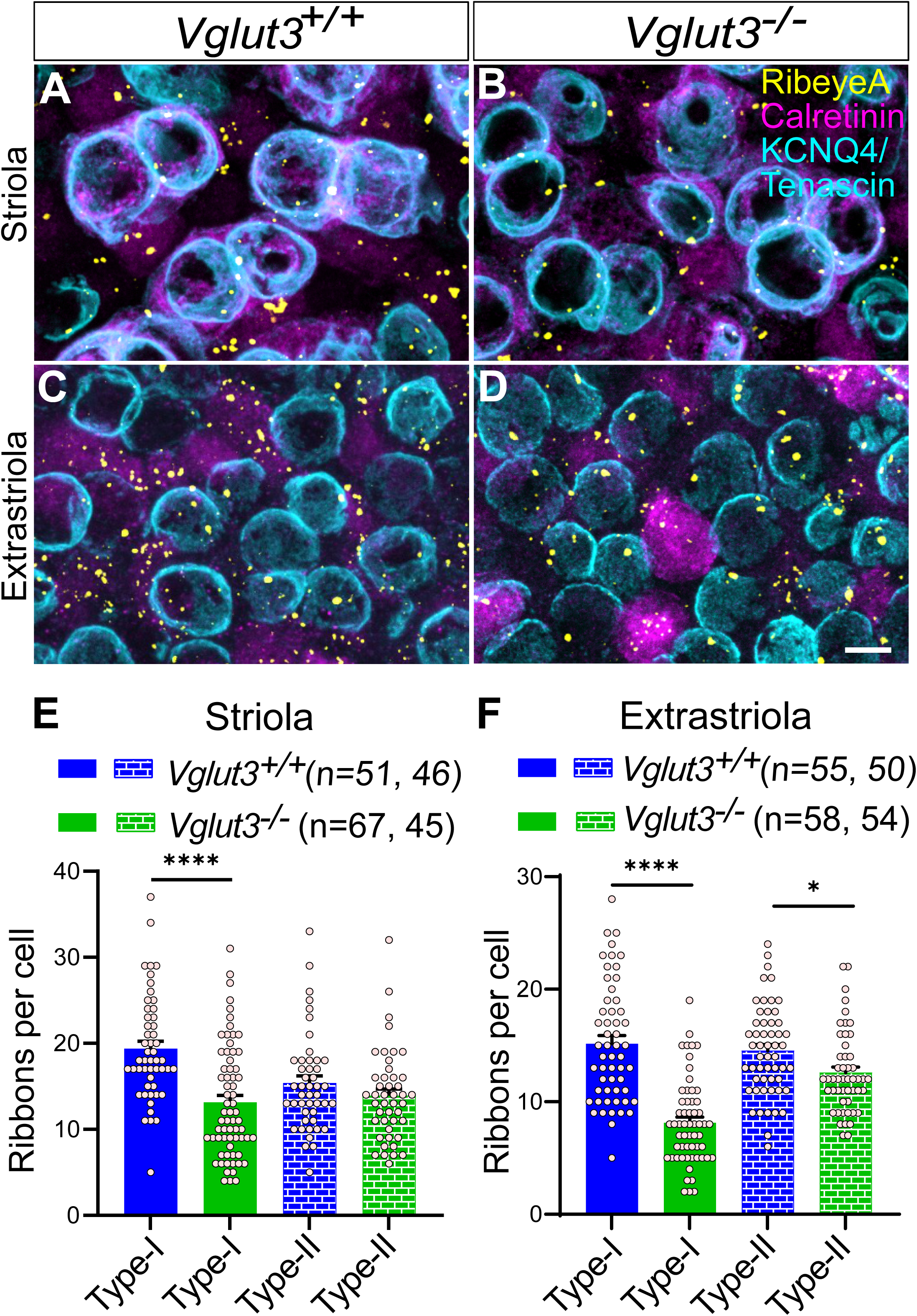
A modest reduction in the abundance of ribbon synapses in the extrastriola of VGLUT3-deficient utriculi. **A-D**) Representative images showing ribbons (RibeyeA) in VHCs of both types. A combination of calretinin, KCNQ4 and Tenascin immunostaining allowed discrimination between type I and type II hair cells in the striola and extrastriola. Scale bar 10 µm. **E-F**) Histograms of ribbons counted in type I and type II hair cells in the *Vglut3^+/+^* versus *Vglut3^-/-^* utricles. Comparable abundance of ribbons was observed in the striola of both VHC types (E). A slight decrease in the number of ribbons per both cell types was detected in the extrastriola of *Vglut3^-/-^* utriculi (F). Student’s *t* test and Holm-Bonferroni correction for multiple comparison, Asterisks in E-F: *p* values from left to right <0.0001, <0.0001 and 0.011, respectively.

## Discussion

In this study, we examined the effects of genetic disruption of VGLUT3 on vestibular function. In the VHCs of VGLUT3-deficient animals, we detected close to zero glutamatergic postsynaptic events and their frequency did not increase upon VHC stimulation. However, the virtually complete loss of quantal synaptic transmission left vestibular sensation predominantly intact, at least, at the level of the behavioral sensorimotor tests. We furthermore observed no upregulation of other VGLUTs nor select ion channels underlying nonquantal transmission. We, therefore, conclude that vestibular endorgans seem not to require classical, quantal synaptic transmission for encoding of vestibular signals. Rather, the nonquantal component of vestibular neurotransmission may transfer sufficient information to support seemingly normal vestibular function.

### Expression profile of VGLUTs in the vestibular endorgans

VGLUT3 has been shown to be pivotal to the development and maturation of the organ of Corti and essential for quantal neurotransmission at the inner hair cell ribbon synapses (Seal *et al*, 2008; Ruel *et al*, 2008; Kim *et al*, 2019). Knocking out *Vglut3* leads to abnormal ribbon morphology in the inner hair cells, profound deafness, and a progressive loss of synapses and spiral ganglion neurons during postnatal development (Seal *et al*, 2008; Ruel *et al*, 2008; Kim *et al*, 2019). Our immunohistochemistry data suggests that VGLUT3 is the primary transporter in the vestibular hair cells as well, in mice. In agreement with previous reports (Zhang *et al*, 2011; Schraven *et al*, 2012), we found strong VGLUT3 expression in type-II VHCs. The expression in type-I VHCs, on the other hand, was either significantly lower or below the threshold for detection, depending on imaging conditions. Interestingly, relatively higher VGLUT3 immunofluorescence intensity was observed in the extrastriolar as compared to striolar type-I VHCs. Since the same immunohistochemistry technique enabled detection of other proteins in the striolar type-I VHCs (see e.g. Fig. 11, 12), we do not ascribe the typical lack of VGLUT3 immunofluorescence in the striolar type-I VHC to factors such as antibody penetrance and thinner cytosol. Therefore, we conclude that VGLUT3 expression in the type-I striolar VHCs indeed must be relatively low, which is also consistent with previous observations (Schraven *et al*, 2012). In this respect, it is also worth noting that the expression of *Vglut3* mRNA seems to be very variable in type I VHCs with some cells showing no to little signal, even at the mRNA level (McInturff *et al*, 2018; see **Suppl. Fig. 2A**). We conclude that while VGLUT3 may function as the primary transporter for packaging glutamate into synaptic vesicles in type-II VHCs and extrastriolar type-I VHCs, striolar type-I VHCs may not depend on VGLUT3. Similarly, the apparently normal sensorimotor and balance behaviors can’t be attributed to the other classical neuronal VGLUTs (VGLUT1 and 2) either, owing to negligible immunofluorescence at the base of the VHCs as well as the scarcity of mRNA spots in the vestibular endorgans, suggesting little to no VHC expression of these transporters.

Nevertheless, in our postsynaptic recordings in *Vglut3^+/+^*animals, which included extrastriolar and striolar calyces, we always observed glutamatergic events. This can have one of the following explanations: 1.) the low levels of VGLUT3, which, in the striola, may fall below the threshold detectable by immunohistochemistry, are sufficient to support quantal neurotransmission at the type-I VHC-calyx synapse, 2.) the recorded events stem from VHC-II-calyx outer leaflet synapses, or 3.) in case of calyces from dimorphic neurons, the recorded events are being sampled from a nearby VHC-II owing to several boutons in a dimorphic afferent which synapse onto them. While we cannot safely exclude one or the other possibility, the outer leaflet synapses have been reported to be relatively scarce (Hamilton, 1968; Gulley & Bagger-Sjoback, 1979; Holt *et al*, 2007) and we thus consider it rather unlikely. Membrane capacitance recordings suggest low levels, if any, of quantal transmission from mature striolar type-I VHCs (Spaiardi *et al*, 2022, 2025), possibly favoring the latter two hypotheses above. VGLUT3 immunofluorescence intensity was ∼6-25 times stronger in mouse utricle VHCs in comparison to VGLUT2, in cells where VGLUT2 was detected. Interestingly, expression of VGLUT1 and VGLUT2 in the utricle and crista was predominantly in the afferent endings and seemed to be complementary, with VGLUT1 primarily localized in the central parts of the vestibular organs (striola in the otolith organs and center in the cristae) and VGLUT2 primarily outside the central region. Their immunofluorescent signals close to the VHC necks co-localized with the calyx marker Na^+^K^+^ ATPase, suggesting these transporters may play some unrecognized role on the synaptic vesicles reported at the putative calyx-to-VHC synapses expressing synapsin I and synaptophysin (Scarfone *et al*, 1988). It is noteworthy; however, there have been no reports about the presence of glutamate receptors at the VHC necks. Further investigation is required to understand the possible roles of VGLUT1/2 in the calyces.

### Glutamatergic quantal synaptic transmission at the VHC ribbon synapses seems to be dispensable for gross vestibular function

Our electrophysiology data shows that knocking out *Vglut3* in mice leads to an abolishment of the excitatory postsynaptic quantal currents, which in the past have been shown to be solely glutamatergic in both VHC types (Sadeghi *et al*, 2014; Holt *et al*, 2006). The apparent absence of VGLUT1/2 or very low expression of VGLUT2 in some VHCs together with the lack of postsynaptic activity in the *Vglut3^-/-^* calyces implies that VHC ribbon synapses in the *Vglut3^-/-^* mice are inactive in the quantal sense, as in cochlear inner hair cells. On the other hand, spontaneous spiking activity in the vestibular neurons *in vivo* seems not to be impaired by abrogation of *Vglut3*. This is in contrast with the data on the *Vglut3^-/-^* auditory afferent fibers, which are inactive when driven by sound or in the absence of sound stimulation (see Fig. 10), and suggests a divergent underlying mechanism. We hypothesize that while in the cochlea, both, spontaneous and stimulus-driven afferent fiber activity depend on vesicular glutamate release (but note that “spontaneous” activity was still detected in the absence of otoferlin; e.g. Pangrsic *et al*, 2010), in the vestibular neurons, both spontaneous and stimulus-driven spikes are present in the absence of glutamatergic transmission at the first vestibular synapse. This is supported by our data of apparently normal spontaneous activity in the vestibular neurons and the sensorimotor tests as a battery of behavioral tests performed on *Vglut3^-/-^*and *Vglut3^+/+^* mice failed to show obvious balance deficits or malfunction, consistent with previous publications (Seal *et al*, 2008; Regalado Núñez *et al*, 2024). What is more, while older *Vglut3^-/-^*mice required on average a longer time to turn on the pole (and swam faster), which we tend to ascribe to increased anxiety (Amilhon *et al*, 2010; Horváth *et al*, 2017), no deficiencies were observed on the rotarod. This latter test is suggested to better probe the potential (mal)function of the vestibular peripheral zones (Regalado Núñez *et al*, 2024; Ono *et al*, 2020). Our data suggest no abnormalities in excitability of the calyces upon current injection. Furthermore, while our dataset on vestibular ganglion neuron spiking is limited, a recent larger dataset supports the conclusion of grossly unaltered excitability of vestibular ganglion neurons (Regalado Núñez *et al*, 2024). Moreover, the biophysical diversity of vestibular ganglion neurons was preserved in the absence of glutamatergic transmission (Regalado Núñez *et al*, 2024).

Our finding of apparently normal vestibular afferent spiking in *Vglut3^-/-^* goes along with prior experiments showing that the blockers of glutamatergic transmission do not prevent spontaneous firing in mammalian calyx terminals (Meredith & Rennie, 2015). Furthermore, the block of mechanotransduction may influence but not abrogate spontaneous firing of vestibular neurons (Horwitz *et al*, 2014; Meredith & Rennie, 2015). Our data is further consistent with reports in other mutants where knocking out crucial exocytosis related genes, such as *Cacna1D* or *Otof*, coding for voltage-gated calcium channel Ca_v_1.3 and otoferlin, from mature VHCs abolished calcium dependent exocytotic vesicular release, yet, left the calyceal postsynaptic action potential firing unaffected (Spaiardi *et al*, 2022; Manca *et al*, 2021). This was also true for calyxes driven by type I VHCs that displayed calcium dependent exocytotic release 10 times smaller than the type II VHCs. It was thus suggested that type II VHCs, due to their large vesicle pools, plausibly support tonic release, while type I VHCs with their restricted pool size putatively support phasic release (Spaiardi *et al*, 2020, 2022).

Is there compensation by another system that allows the animals to function normally in the absence of VGLUT3? Non-quantal neurotransmission has gained immense traction in the past few decades, proving to be a crucial part of the communication network between the peripheral vestibular system and the central nervous system. It is hypothesized that as calyces evolved from boutons and proto-calyces, selective pressure preferred the presence of direct resistive coupling between the type I VHCs and calyces that led to ultrafast signal transmission (Contini *et al*, 2022). A recent computational model demonstrated the importance of calyceal cleft electrical potential and cleft K^+^ in modulating the gain and phase of non-quantal transmission (Govindaraju *et al*, 2023). This study also showed that a dynamic cleft potential reduces transmission latency in such structures. Furthermore, calyx morphology, such as calyx height, was found to be a determinant for the latency of action potential, with longer calyces having lesser latency of transmission. All of the above are in line with the idea that calyces evolved to support non-quantal transmission and to speed up phasic vestibular signaling for driving very fast vestibular reflexes (Govindaraju *et al*, 2023). We thus looked at the various components of the non-quantal mechanisms.

### The absence of glutamatergic synaptic transmission affects various ion channels in VHCs and calyces

We observed a shift of approximately −8 mV in the HCN2 activation curves towards more negative half-activation potentials in the utricular calyces of *Vglut3^-/-^* mice. While such a hyperpolarized shift, at first glance, seems counterintuitive (as described below), as HCN channels are known to sharpen post synaptic response and increase membrane excitability by reducing membrane resistance at negative potentials (Meredith *et al*, 2012), we argue that this could be an indicator of developmental changes that favor more excitable VGNs (Ventura & Kalluri, 2019). An upregulation of HCN channel currents during development leads to more recruitment of low-voltage-activated currents, which counterintuitively reduce vestibular ganglion neuron excitability, turning them into phasic neurons (Ventura & Kalluri, 2019). Hence, a hyperpolarized shift in the activation curve implies a lower I_h_ which may contribute to lower expression of K_V_ channels or low I_kl_, which in turn would lead to a higher population of low threshold tonic firing neurons. Cells lacking VGLUT3, and therefore quantal transmission, might favor such a mechanism to augment non-quantal neurotransmission. Interestingly, a recent study suggested possibly reduced I_kl_ in large, putative central-zone vestibular ganglion neurons in *Vglut3^-/-^* mice, but not in those innervating the periphery (Regalado Núñez *et al*, 2024). Our immunohistochemical analysis of KCNQ4 signal did not reveal significant differences in the striola, but a trend toward lower levels was detected. We acknowledge that the immunohistochemical analysis has limited sensitivity and does not reveal the biophysical properties of the ion channels as electrophysiology does, nevertheless, the observation of lower KCNQ4 expression in the extrastriola is intriguing and requires further examination.

Our immunohistochemistry data further suggests a reduction in the HCN1 levels in both the striolar and extrastriolar regions in the *Vglut3^-/-^*. There is a dramatic increase in HCN1 expression in the VHCs (I and II) in the first postnatal week, which then levels off and stabilizes (Rüsch *et al*, 1998; Horwitz *et al*, 2011). Since our immunohistochemistry was performed at mature ages, we do not ascribe the observed changes to a putative lag in development, but rather a downregulation of the channel. Calyces show limited HCN1 expression (Horwitz *et al*, 2014, 2011). Limited spatial resolution hindered a clear distinction between the post- and presynaptic membranes, but due to a significantly more prominent expression of HCN1 in the VHCs, we presume that the observed differences in the expression levels in the two genotypes are most likely related to the changes in the VHCs. However, our data is compatible with the idea of a reduction in both, VHCs and calyces. In this respect, a putative loss of HCN1 in the calyces could underlie the negative activation shift of the calyceal HCN currents (Horwitz *et al*, 2014), as observed here. Reduced inward calyceal HCN2 current and lesser HCN1 current into hair cells could potentially add more positive ions to the cleft. This could in turn alter the cleft electrical potential and consequentially the driving force for cations. This could then facilitate depolarization of both the pre and postsynapse. Another possible explanation for the reduction in I_h_ may be an energy/resource saving mechanism deployed by the cells in the absence of quantal transmission. It is speculative to comment on the possible causation or correlation of the activity/levels of VGLUT3, HCN and KCNQ channels at this point. Whether one or more of the mechanisms mentioned above are working simultaneously or independently or in a well-coordinated sequential manner in microdomains is a matter for further investigation.

In summary, even though transmission at the VHC synapse is glutamatergic and VGLUT3 is the main vesicular glutamate transporter, abrogating VGLUT3 seems to be inconsequential for vestibular sensation. This has also been suggested by a study applying AMPA receptor antagonists which showed important evidence that synchronized vestibular compound action potentials are driven by fast non-quantal transmission (Pastras *et al*, 2023). This further supports the idea that non-quantal synaptic transmission is plausibly sufficient in transmitting enough information to support vestibular function, although milder vestibular defects, not detected by our methods, cannot be entirely excluded. While nonquantal transmission has been observed elsewhere (e.g. Blot & Barbour, 2014; Chiang *et al*, 2019; Han *et al*, 2020), such a prominent role has not been ascribed to it yet. In this respect, vestibular synaptic transmission may have evolved a unique way of encoding sensory information to enable more rapid transmission to serve fast reflexes (Eatock, 2018).

## Materials and Methods

### Animals

*Vglut3* knockout mice (*Vglut3^-/-^*) on a C57BL/6J (B6) background were obtained from The Jackson Laboratory (RRID: IMSR_JAX:000664). Herein, *Vglut3^-/-^* and *Vglut3^+/+^*mice are sometimes referred to as KO and WT, respectively. *Vglut3* floxed mice (*Vglut3^fl/fl^*) on a B6 background were obtained from Rebecca P Seal. *Vglut3^fl/fl^*were driven by CAGGCreERTM mice obtained from The Jackson Laboratory (RRID:IMSR_JAX:004682). Here, Cre+ mice are sometimes referred to as *Vglut3*-conditional KO. All experiments complied with national animal care guidelines and were approved by the University of Göttingen Board for animal welfare, the animal welfare office of the state of Lower Saxony, and the animal studies committee at Washington University in St. Louis.

### *In vitro* electrophysiology

Mice were anaesthetized using CO_2_ administration at the rate of 8-10 ml/min. The utricles were dissected out in external solution (recipe below) and treated with 100 µg/ml protease XXIV for 5-7 mins, followed by removal of the otolithic membrane by an eyelash glued to a Q-tip. Isolated utricles were then positioned under nylon threads attached to a grid in a recording chamber containing the extracellular solution that was perfused through the chamber at a rate of ∼0.4 ml/min. Cells were visualized with a 60X water immersion lens on an upright microscope from Olympus. Whole-cell currents were recorded from the calyx afferent endings in the striolar and extrastriolar regions of utricles belonging to P17-25 mice of either sex. The intracellular solution for the afferent recordings contained the following (in mM): 20 KCl, 110 K-methanesulfonate, 0.1 CaCl_2_, 5 EGTA, 5 HEPES, 5 Na_2_ phosphocreatine, 4 MgATP, 0.3 Tris-GTP, 290 mOsm, pH 7.2 (KOH). The extracellular solution had the following composition (in mM): 5.8 KCl, 144 NaCl, 0.9 MgCl_2_, 1.3 CaCl_2_, 0.7 NaH_2_PO_4_, 5.6 glucose, 10 HEPES, 300 mOsm, pH 7.4 (NaOH). Equimolar NaCl was replaced with KCl for some experiments to increase the extracellular K^+^ concentration to 22 or 40 mM. Unless stated differently, all chemicals were purchased from Sigma-Aldrich. Na-channel blocker tetrodotoxin (TTX) was added in the extracellular solution at 500 nM to block action currents. Alexa 568 hydrazide (ThermoFisher Scientific, #A10437) was added to the internal solution at 100 µM to visualize the patched calyces. Supporting cells around the calyx-of-interest were gently cleaned with a wide-mouth pipette, followed by positive pressure with a patch pipette to clear out debris. Predominantly, complex calyces in striolar or extrastriolar regions were targeted. Patch pipettes with resistance ranging from 6-10 MΩ were pulled from borosilicate glass capillaries with filament (GB150F, 0.86 × 1.50 × 80 mm; Science Products) using P1000 micropipette puller (Sutter Instruments Co). Recordings were performed at room temperature and were acquired using a HEKA EPC10 amplifier and Patchmaster software. Calyces were clamped at −79 or −103 mV. Liquid junction potential of −9 mV was corrected off-line for the recordings. Data were sampled at 25 µs and filtered at 7.3 kHz. Membrane capacitance and series resistance were calculated by the HEKA software. Series resistance was not compensated. The average holding currents at −79 mV were −91 ± 29 pA (*Vglut3^+/+^*) and −182 ± 56 pA (*Vglut3^-/-^*), which is in line with similar recordings from rat cristae (Sadeghi *et al*, 2014). As previously shown, the larger holding currents are indicative of open ion channels at hyperpolarizing potentials rather than unspecific leak. Since unspecific leaks cause cells to be depolarized, we corroborate these data with the membrane potentials measured intracellularly. The average resting potentials were −70.2 ± 1.3 mV (*Vglut3^+/+^*) and −71.5 ± 2.6 mV (*Vglut3^-/-^*), which were indicative of healthy seals. Cells with holding currents larger than 350 pA at −79 mV were discarded. However, it is to be noted that the *Vglut3^-/-^* showed a trend (not significant) towards slightly larger holding current than the *Vglut3^+/+^*. The I_h_ recordings of currents through the HCN channels were done immediately after breakthrough to the whole-cell configuration to avoid potential run down of currents that can occur quickly. Here, the cells were hyperpolarized from rest to −159 mV for 650 ms in 10 mV steps. Tail currents were measured at 40 ms after the return of the voltage to the resting value of −79 mV, close to the reversal potential of the K-channels and below the activation range of the voltage-gated Ca- and Na-channels. Onset current (I_ons_) was measured at the peak (occurring at ∼65 ms) and I_end_ was the steady state current measured at the end of the voltage step.

Excitatory postsynaptic currents (EPSCs) were detected using the NeuroMatic software tool (Rothman & Silver, 2018). Further analysis was done using custom-written routines in Igor Pro (Wavemetrics). Postsynaptic current decays were fitted using a single or double exponential (slow decay time constant at least 5 times the fast decay). EPSC decay, amplitude and frequency were analyzed and plotted using customized routines written in Igor Pro and Prism (GraphPad). Overlapping or summated events were classified as complex events, and amplitude and decay analyses were not performed for such events.

### Immunofluorescence and Confocal microscopy

#### Ion channel, synaptic and VGLUT labelling in wholemounts

The inner ears were dissected out from P15-39 *Vglut3^-/-^*, *Vglut3^+/+^* littermates, and WT B6 mice. A small hole was made with forceps in the temporal bone just above the utricle to uncover the pigmented membrane underneath, which was then carefully cut open to expose the utricle. The inner ear was fixed in 4% formaldehyde on ice for ∼45 mins, briefly (1 min) rinsed in deionized water, decalcified by TBD-1 (∼30 s; Shandon TBD-1TM Rapid Decalcifier, #6764001), and washed again in deionized water (1 min). The utricles were then gently excised from the inner ear and transferred to phosphate buffer saline (PBS: Sigma #P4417) for washing. The whole organs were then treated with goat serum blocking and permeabilization buffer solution (16% goat serum, 450 mM NaCl, 0.3% Triton X-100, and 20 mM phosphate buffer, at pH 7.4) for 1h at room temperature, then incubated in a primary antibody concoction in buffer solution overnight at 4°C. The next day they were rinsed in wash buffer (0.3% Triton X-100 in PBS) and incubated in a concoction of secondary antibodies in buffer solution for 1 hour in a dark wet chamber, rinsed again with wash buffer and whole-mounted with mowiol mounting medium on microscopy slides. The following antibodies and dilutions were used: chicken-anti-calretinin (Synaptic Systems, #214106), rabbit-anti-RIBEYE (Synaptic Systems, #192103), mouse-anti-beta-III-tubulin (Biolegend, #801202), rabbit-anti-VGLUT3 (Synaptic Systems, #135203), rabbit-anti-KCNQ4 (Merck, #HPA018305), rabbit-anti-KCNQ5 (Merck, #ABN1372), rabbit-anti-HCN1 (Invitrogen, #PA5-78675), guinea pig-anti-RIBEYE (Synaptic Systems, #192003), rabbit-anti-tenascin (Millipore, #AB19013). The secondary antibodies used were: Alexa Fluor 488 labeled goat-anti-chicken (Invitrogen, #A11039), Alexa Fluor 568 labeled goat-anti-mouse (Invitrogen, #A21134), Alexa Fluor 647 goat-anti-rabbit (Invitrogen, #A21244), Alexa Fluor 647 labeled goat-anti-guinea pig (Invitrogen, #A21450). All secondary and primary antibodies were used at a dilution of 1:200. Confocal images were acquired using an Abberior Instruments Expert Line microscope equipped with 488, 561 and 633 nm excitation lasers, and a 100X oil immersion objective (1.4 NA, Olympus). Images of organs from animals of different genotypes were acquired using identical microscope settings, analyzed and processed in parallel in Fiji/ImageJ software or Imaris (Oxford Instruments), and assembled in Affinity Designer for display. For VGLUT1 and 2, preparation, immunolabeling and imaging of the wholemount utricles were performed as described (Warchol *et al*, 2019). The organs were labeled with DAPI (Thermo Fisher), ßIII-tubulin (Tuj1; Biolegend, #801202), and rabbit anti-Vglut1 (RRID:AB_887875) or rabbit anti-Vglut2 (RRID:AB_887883). Quantification of ribbon synapses was performed in Imaris (Oxford Instruments), using the annotation tool to select the cells of interest and the spots function to detect the ribbons. The XY-diameter of the spots was set to 0.4 µm throughout the analysis and the spot threshold limit to default mode.

#### Hair cell and nerve terminal labelling

Calyx nerve terminals were counted using confocal images obtained from standardized regions of whole-mount utricles (Warchol *et al*, 2019). A Zeiss LSM700 confocal microscope (Carl Zeiss, Germany) was used to acquire images from the medial extrastriolar and striolar regions of individual specimens. Confocal volumes were acquired with 12-bit images with a 63X 1.4 NA oil objective lens with a Z-step of 0.37 µm and pixel size of 50 nm in X and Y, and a pinhole setting of 1 AU. Image stacks were reconstructed and visualized using Volocity software (PerkinElmer, Akron, OH), and cell quantification was conducted from 50 × 50 μm regions of z-slices. Hair cells were labelled with Myosin7a (RRID AB_10015251). Calyces, labelled with anti-ßIII-tubulin (RRID AB_2721321), were counted only if they appeared to fully enclose the basal to mid-regions of a target hair cell at the level of the cell nucleus.

#### VGLUT1-3 labelling in cryosections

For labelling of VGLUT1, VGLUT2, and VGLUT3 in utricules and cristae on cryosections, temporal bones were decalcified in 10% EDTA in PB, cryo-protected in a sucrose series (10, 20, and 30%), frozen in Tissue Freezing Medium (Electron Microscopy Sciences), then cryo-sectioned at 20 µm thickness along the anterior-posterior axis. After rehydration in PBS, sections were permeabilized and non-specific reactivity was blocked with horse serum blocking and permeabilization buffer solution (16% horse serum, 450 mM NaCl, 0.3% Triton X-100, and 20 mM phosphate buffer, at pH 7.4) for 2 hours at room temperature. The rest of the rinsing and staining protocol was identical to the whole-organ method above. Hair cells were labelled with mouse IgG2a anti-Myosin7a (RRID:AB_2148626). VGLUTs were labelled with rabbit anti-Vglut1 (RRID:AB_887875), rabbit anti-Vglut2 (RRID:AB_887883), or goat anti-Vglut3 (RRID:AB_2187701). Airyscan volumes from the center of the utricle were acquired with 16-bit images on a Zeiss LSM 880 Confocal with a 100X 1.5 NA oil objective lens with a Z-step of 0.168 µm and pixel size of 26 nm in X and Y, and a pinhole setting of 1.5 AU. Low magnification images were made with a 20X objective on a Zeiss LSM700.

#### RuneScape

For estimating mRNA levels of *Vglut1* and *Vglut2*, cryosections of three *Vglut3^-/-^* and three *Vglut3^+/+^*3-month-old mice were utilized. Cryosections were prepared as described above and cut at 20 µm thickness along the anterior-posterior axis of the utricle, also containing the posterior crista and vestibular ganglion. RNAscope Multiplex Fluorescent Reagent Kit version 2 (catalog #323110) was used to perform in situ hybridization (ISH) with RNAscope probes according to the protocol provided by the manufacturer for fixed frozen tissue with some modifications (Ferreira *et al*, 2022). Probes for *Vglut1* (Mm-Slc17a7, catalog #416631-C3) or *Vglut2* (MmSlc17a6, catalog #319171-C2) were visualized with Cy5 fluorescence (NEL745001KT-TSA Plus Cyanine 5, Akoya Biosciences) fluorophore on separate slides.

After the ISH protocol, slides were blocked with 10% normal donkey serum in 1x tris buffered saline (TBS) and 1% bovine serum albumin (BSA) solution for 1 hour at room temperature (RT). All sides were then incubated with primary antibodies in 1x TBS+ 1% BSA-Myosin7a (RRID AB_10015251) and a combination of ßIII-tubulin, (RRID AB_2721321) and Neurofilament H (RRID:AB_1005601) overnight at 4°C, then washed and stained with secondary antibodies for 2 hours at RT – Alexa Fluor 594 labeled donkey-anti-mouse (Jackson Immuno, 715-586-150, 1:300) and Alexa Fluor 488 labeled donkey-anti Rabbit (Jackson Immuno, 711-545-152, 1:200). Samples were washed and mounted using ProLong Gold Antifade with DAPI (Thermo Fisher Scientific, catalog #P36931). Images were collected on the Zeiss LSM 700 as described above. Immunofluorescent spots in vestibular neurons were counted using the spots function in Imaris as described above (ribbon counting). The criterion for spot size, based on the XY spot diameter was set to 0.1 or 0.2 µm.

### Behavior tests

#### Rotarod

The rotarod protocol included three conditions: a stationary rod, a rod that rotated at a constant speed (2.5 rpm for 60 s maximum), and a rod that rotated at an accelerating speed (2.5 – 10.5 rpm over 1– 180 s maximum). The protocol consisted of three training sessions with each session including one stationary rod trial, two constant speed rotarod trials, and two accelerating speed rotarod trials. Sessions were separated by 4 days to minimize motor learning and time spent on the rod was used as the dependent variable to assess performance. All trials were performed using an Economex Rotarod (Columbus Instruments, Columbus, OH). WT and KO mice were tested at 3 and 8 months of age.

#### Open Field Circling

Circling behavior was measured in an open field apparatus consisting of a transparent (16”16”x15”) polystyrene enclosure, surrounded by a frame containing a 16×16 matrix of photocell pairs. Mice were habituated to the room for 30 min prior to testing, then placed into the chamber and behavior recorded for 30 min. A camera was mounted above the apparatus and clockwise and counterclockwise rotations were recorded and measured using the ANYMaze software (Stoelting, Inc). WT and KO mice were tested at 3 and 8 months of age.

#### Righting reflex

In the surface righting reflex test, mice were placed onto their backs in an empty cage lined with a plastic bench pad and the time taken to flip over into the normal position with respect to gravity was recorded, up to 60 s, and the mean of 3 consecutive trials was reported as the dependent variable (McCullough *et al*, 2024). In the air righting reflex test, mice were observed and scored for their ability to right themselves completely from an inverted position in freefall from a height of 50 cm onto a padded surface. Each mouse was tested in five consecutive trials and received a score indicating if it landed on all four paws or did not land on all four paws. WT and KO mice were tested at P5 and P14 for surface righting reflex and at P18 for air righting reflex. *Vglut3*-conditional KO (Cre+ and Cre-) were tested at 4 months of age on both reflex tests after receiving tamoxifen at 3 months, and after confirming deafness at 3.5 months.

#### Sensorimotor battery

Walking initiation, ledge, platform, and pole tests were used to evaluate sensorimotor function. Test duration was 60s, except for the pole test, which was extended to 120s. For walking initiation, time for an animal to leave a 21×21cm square on a flat surface was recorded. For ledge and platform tests, the time the animal was able to balance on an acrylic ledge (0.75cm wide and 30cm high), and on a wooden platform (1.0cm thick, 3.0cm in diameter and elevated 47cm) was recorded, respectively. The pole test was used to evaluate fine motor coordination and balance by quantifying time to turn 180° and climb down a vertical pole. Time in each task was manually recorded and the average of two trials on each test was used for analyses. WT and KO mice were tested at 3 and 8 months of age.

#### Inclined and inverted screens

The screen tests assess a combination of coordination and strength. The mouse was placed head oriented downward in the middle of a mesh wire grid measuring 16 squares per 10 cm, elevated 47 cm and inclined to 60°, 90°, or inverted. Time taken to turn upward 180° and climb to the top of the screen was recorded up to 60 s, and the mean of two 60 s trials was reported as the dependent variable. WT and KO mice were tested at 3 and 8 months of age.

#### Acoustic startle

The acoustic startle reflex was assessed by recording the mouse’s whole-body flinch in response to a white-noise acoustic stimulus of varying intensity using Kinder Scientific Startle Reflex equipment and software (Kinder Scientific, Inc). Mice were placed on a platform in an enclosed chamber to record movement with a piezoelectric force transducer under the platform. Auditory function was assessed by measuring behavioral responses to auditory stimuli pulses (40 ms broadband burst) presented at 80dB, 90dB, 100dB, 110dB, and 120dB above background noise (65dB), as well as no-stimulus trials. Each trial type was presented 10 times in a random order over a 20 min session, with inter-trial-intervals ranging from 5-15 s between trials. The average force response to each sound pressure level was analyzed as a measure of auditory function. WT and KO mice were tested at 3, 8, and 12 months of age.

#### Swim test

To test vestibular function and swimming ability, mice were placed in a pool (114cm diameter) of room temperature water and assessed for three 60 s trials. Swimming ability was recorded using an overhead camera and analyzed with the ANYMaze software program (Stoelting, Inc). Following each 60 s trial, mice were placed in warming cages to dry and given 30 min rest between trials. Total distance traveled and swimming speed were analyzed as measures of swimming ability. WT and KO mice were tested at 3 and 8 months of age.

### Single Unit Recordings

All single unit recordings of the VIII^th^ nerve were from the left ear. Mice were anesthetized with Ketamine/Xylazine cocktail (80/15 mg/kg, supplemented with half-doses as needed) and placed dorsal side up in a stereotaxic frame in such a way that the head was level side-to-side, but tipped slightly down posteriorly. Temperature was maintained near 37 ⁰C using a thermostatic probe and heating pad (Fredrick Haer, FHC). The skin was removed from the entire top of the cranium and the underlying bone was scraped and dried to accommodate marking of the recording site and gluing of a small metal plate across the head, from just behind the eyes to Bregma using Super Glue. The plate was affixed in the frame and a ∼2×2 mm plate of bone was removed from a region spanning from Bregma to the lambdoid suture and beginning 1 mm peripheral from the midline toward the left ear. The top ∼2 mm of cerebellum was then removed by suction and the exposed brain was covered in mineral oil. Recording electrodes (10-50 MΩ) were pulled from borosilicate capillary glass using a Sutter Model P-97 and filled with 0.9% NaCl. The microelectrode medium was chosen so as not to flood the recording site with high levels of Na^+^ or K^+^ in cases of electrode breakage. Electrodes were aimed using a combination of quasi-stereotaxic coordinates and landmarks, and driven into the brain in 5 µm steps using a Kopf 650 hydraulic micropositioner. Vestibular neurons, which also make up the cochlear nerve with cochlear neurons, were often encountered and identified as such by a) close proximity to acoustically-driven neurons, b) lack of acoustic activity, and c) the frequent presence of regular spiking activity.

Sound stimuli were presented in near-field using a TECTONIC TBM54C30-8/ TDT ES-1 hybrid speaker and calibrated offline by placing a BK ¼ inch microphone where the pinna would be. ABR thresholds to clicks (100 µs, 1000x, 20/s) were obtained prior to recording and rechecked periodically. Click responses were recorded using Grass needle electrodes in combination with TDT hardware and BiosigRZ software. Single unit microelectrodes signals were directed to a Warner Instruments Model IE 210 intracellular amplifier, and spikes were detected using a peak discriminator (FHC). While tracking for auditory units, 70-90 dB SPL non-stationary noise bursts (5/s, 100 ms, 0.5 – 80 kHz) were presented as search stimuli. Tone presentation and processing of single unit firing rate and timing utilized TDT RZ6 hardware and custom Labview routines. For each unit isolated, data collected included frequency tuning curves (2 - 78.7 kHz, log frequency steps), input/output curves at the identified characteristic frequency (CF), and post-stimulus-time histograms (PSTHs) at CF (90 dB SPL, 100x, 100 ms tonebursts). Spontaneous firing rates (SR) for each unit were calculated as the average rate measured during four 5 s intervals placed between stimulus runs. Following Taberner and Liberman (2005), joint analysis of electrode depth, coefficient of variation of interspike intervals, and first-spike latency was used to distinguish cochlear neurons from cochlear nucleus neurons. Our tuning curve algorithm was as follows: Stimulus levels above 90 dB SPL were not permitted. ‘Response’ decisions required that more spikes occur during a 100 ms stimulus window than in a 100 ms no-stimulus window positioned 100 ms later. The criterion spike rate increase was +10% relative to the SR estimate from the previous 5s counting period. Frequencies were presented in low-to-high order. In the tail of the tuning curve, the frequency was advanced using a stimulus level of 90 dB SPL until a response was encountered. The stimulus level was then lowered in 5 dB steps until there was no response, then the frequency was advanced until a response was elicited. The tip of the tuning curve was recognized as the first descending stimulus level where increasing frequency steps elicited no response. At that point, the level was increased by 10 dB and the same frequency range was again scanned at progressively lower sound levels in 1 dB steps. Above CF, the sound level increased in 5 dB steps until there was a response, then the frequency was incremented and the process was repeated.

### Video-EEG

VGLUT3 KO and wild-type control mice underwent continuous video-EEG monitoring starting at 6 months of age for two weeks, using established methods for implanting epidural electrodes and performing continuous video-EEG recordings, as described previously (Erbayat-Altay *et al*, 2007; Zeng *et al*, 2008). Briefly, mice were anesthetized with isoflurane and placed in a stereotaxic frame. Epidural screw electrodes were surgically implanted and secured using dental cement for long term EEG recordings. Five electrodes were placed on the skull: one right and one left central (parietal) electrodes (2 mm lateral to midline, 2 mm posterior to bregma), two frontal electrode (0.5 mm anterior and 0.5 mm to the right and left of bregma) and one occipital electrode (1 mm posterior and 0.5 mm to the right or left lambda). The typical recording montage involved two EEG channels with the right and left central “active” electrodes being compared to either the frontal or occipital “reference” electrode. Video and EEG data were acquired simultaneously with Stellate video-EEG system. Continuous 24/7 video-EEG data were obtained for two weeks and analyzed by a blinded reviewer for spike and wave discharges or electrographic seizures (rhythmic spike discharges that evolved in frequency and amplitude lasting at least 10 s, or bursts of spike and wave discharges associated with a behavioral change on video).

## Statistical analysis

Statistical significances between two groups were calculated with Student’s *t* test if the data were distributed normally or Mann-Whitney *U* test if the data were not normally distributed. Normality of data was tested using Jarque-Berra or Kolmogorov-Smirnov test and variances were compared using *F*-test. Data in figures are reported as mean±SEM unless indicated differently. For comparisons between more than two conditions, one-way ANOVA along with Tukey’s HSD *post hoc* or Holm-Bonferroni tests were utilized. In the behavioral tests, Chi-Square test, paired *t*-tests, or 2-way repeated measures ANOVA with Bonferroni corrected multiple comparisons were used. Asterisks represent the following: \**p* < 0.05, \*\**p* < 0.01, \*\*\**p* < 0.001, and \*\*\*\**p* < 0.0001.

## Author contributions

MM, TP, MEW, RPS, and MAR designed the study and secured funding. MM performed patch-clamp experiments and immunohistochemistry with respective analysis. RM performed immunohistochemistry and analysis of RNAscope data. TP contributed to analysis of patch-clamp data. AYH performed single-unit recordings from the vestibular nerve. KKO and VM performed single-unit recordings from the auditory nerve. MX and ZS processed tissues for anatomical studies and did confocal microscopy. NR and MW provided electro-encephalogram. SJL and RPS performed RNAscope labelling. MEW performed immunofluorescence, microscopy, and analysis. SEM and CMY provided and analyzed behavioral data. MAR performed immunofluorescence, microscopy, and analysis of confocal images and single-unit recordings. MM, TP, RM and MAR wrote the manuscript. All authors edited the manuscript.

## Acknowledgments

The research was supported by NIH grant R01DC014712 (MAR), R01DC006283 (MEW), NS082650 (RPS), and ProfProgramIII 01FP19073K and MWK 22-76251-99-17/19 (TP). Research reported in this publication was further supported by the Eunice Kennedy Shriver National Institute Of Child Health & Human Development of the National Institutes of Health under Award Number P50 HD103525 to the Intellectual and Developmental Disabilities Research Center at Washington University and the Institute of Clinical and Translational Sciences at Washington University (NIH CTSA Grant #UL1 TR002345). We would like to thank Mrs. Jamie Hicks, Mr. Evan Daniels-Day, and Mr. Cory Cearlock for help with behavioral data collection, and Ina Preuss for genotyping. We would also like to thank Matthew Kelley for *Vglut1* and *2* mRNA expression data in Supplemental Figure 2A that is available at gEAR (https://umgear.org/).

